# Organizational principles governing synapse types in a whole-brain connectome

**DOI:** 10.64898/2026.06.23.733969

**Authors:** Amit Gross, Majd Farah, Subhajit Jana, Sven Dorkenwald, Davi Bock, David Deutsch

## Abstract

Neurons are classically viewed as polarized elements in which dendrites receive synaptic input and axons provide output. However, many neurons contain intermingled presynaptic and postsynaptic sites, enabling non-canonical forms of connectivity beyond axo-dendritic interactions. How such mixed polarity shapes connectivity across an entire nervous system remains unknown.

Here, using the complete *Drosophila* connectome of an adult brain (approximately 140,000 neurons and >80 million synapses), we systematically mapped synapse types across neuronal compartments, cell types, and brain regions. We found that mixed polarity is widespread yet highly structured across neuronal populations. Approximately one third of synapses between polarized neurons are non-canonical, including axo-axonic, dendrodendritic, and dendro-axonic interactions. Remarkably, synapse type can be predicted from a simple axis of neuronal complexity derived from morphology and connectivity. Low-complexity neurons, which dominate early visual circuits, preferentially interact through dendrites, whereas larger and more highly connected neurons enriched in central brain regions preferentially interact through axons. Mixed polarity also strongly promotes reciprocal connectivity, which is dominated by non-canonical motifs. Finally, we identified postsynaptic terminals on presynaptic boutons as a widespread anatomical motif associated with local reciprocal and non-canonical interactions. Together, these findings reveal organizational principles linking neuronal architecture, synapse type, and circuit connectivity across a complete brain.

## Introduction

Neurons are classically viewed as polarized cells in which information flows from dendrites, the primary recipients of synaptic input, to axons, the principal sites of synaptic output^1–5^. This principle of dynamic polarization, first articulated by Ramón y Cajal, has provided a remarkably successful framework for understanding neural circuit organization. In many systems, dendritic and axonal compartments are indeed anatomically distinct and support largely segregated input and output functions^5–10^. However, numerous studies across vertebrate and invertebrate nervous systems have revealed neurons in which presynaptic and postsynaptic sites coexist within the same arbor, branch, or even subcellular compartment^5,9–19^. Such “mixed polarity” enables synaptic interactions that fall outside the canonical axon-to-dendrite (AD) configuration, including dendrite-to-dendrite (DD), axon-to-axon (AA), and dendrite-to-axon (DA) communication^10,12–14,19–30^.

These non-canonical interactions have been reported in a wide range of circuits. dendrodendritic synapses are prominent in the mammalian olfactory bulb, where they support reciprocal inhibition and gain control^19,29^. axo-axonic interactions regulate transmitter release in diverse vertebrate circuits and can profoundly influence information flow^10–12,26,31,32^. In insects, mixed pre- and post-synaptic specializations have been described in multiple circuits, including the mushroom body, central complex, optic lobe, and antennal lobe^14,15,23,24,33–39^. Collectively, these observations suggest that neurons often interact through a richer repertoire of synaptic architectures than the canonical dendrite-to-axon framework implies. Yet despite more than a century of observations, we still lack a quantitative understanding of how mixed polarity is distributed across a complete nervous system and whether general principles govern the use of different synaptic interaction types.

The emergence of dense connectomic reconstructions has created an opportunity to address these questions systematically. Previous work introduced the Segregation Index (SI), an entropy-based measure of neuronal polarity that quantifies the degree to which presynaptic and postsynaptic sites segregate into axonal and dendritic compartments^40^. SI has been applied to individual circuits and to the complete first-instar larval *Drosophila* nervous system^41^, revealing substantial variability in neuronal polarity^40,41^. More recently, whole-brain connectomes of the adult fly have become available, including FlyWire, which provides synapse-resolved reconstructions for essentially all neurons in the adult female brain^42–44^. These resources make it possible to move beyond individual examples and ask whether mixed polarity follows reproducible organizational rules at the scale of an entire brain.

Several fundamental questions remain unresolved. Is mixed polarity simply a consequence of compact brain architecture, developmental constraints, or reconstruction artifacts, or does it reflect functionally meaningful circuit organization? Are non-canonical synapse types distributed randomly across neurons and brain regions, or can their occurrence be predicted from neuronal morphology, connectivity, and circuit position? How does mixed polarity shape reciprocal connectivity, a ubiquitous feature of neural networks^45–52^? Finally, are there common anatomical motifs that support local interactions between input and output compartments?

Here, we combine whole-brain connectomics, polarity analysis, and synapse-level classification to generate a brain-wide map of synaptic interaction types in the adult *Drosophila* brain. We show that mixed polarity is widespread yet highly reproducible across cell types and hemispheres. We find that the use of canonical and non-canonical synapse types is strongly structured by neuronal identity and by a simple axis of neuronal complexity derived from morphology and connectivity. These organizational principles predict the prevalence of axo-axonic, dendrodendritic, and dendro-axonic interactions across brain regions and reveal that reciprocal connectivity is dominated by non-canonical motifs. Finally, we identify postsynaptic terminals located on multisynapse boutons (MSBs) as a widespread anatomical substrate for local reciprocal and non-canonical interactions. Together, our results suggest that mixed polarity is not merely a deviation from canonical neuronal organization but a fundamental component of how neural circuits are constructed and interconnected.

## Results

### Mixed polarity is widespread across the adult brain connectome

To quantify the distribution of presynaptic (output) and postsynaptic (input) terminals across all neurons, we used a new synapse annotation resource^53^ that improves upon the synapse detections underlying the FlyWire/FAFB (full adult fly brain) connectome^54^ (Fig. 1A). The degree of segregation between presynaptic and postsynaptic terminals varies widely across neurons. In some cells, presynaptic and postsynaptic sites occupy largely distinct domains (Fig. 1B, C), whereas in others they are extensively intermingled throughout the neuronal arbor (Fig. 1D). We refer to this intermingling of presynaptic and postsynaptic terminals on the same neuronal branches as “mixed polarity.” In neurons with clear segregation, input and output domains (dendrites and axons, respectively) can be identified directly from the spatial distribution of synapses (Fig. 1C).

**Fig. 1:**
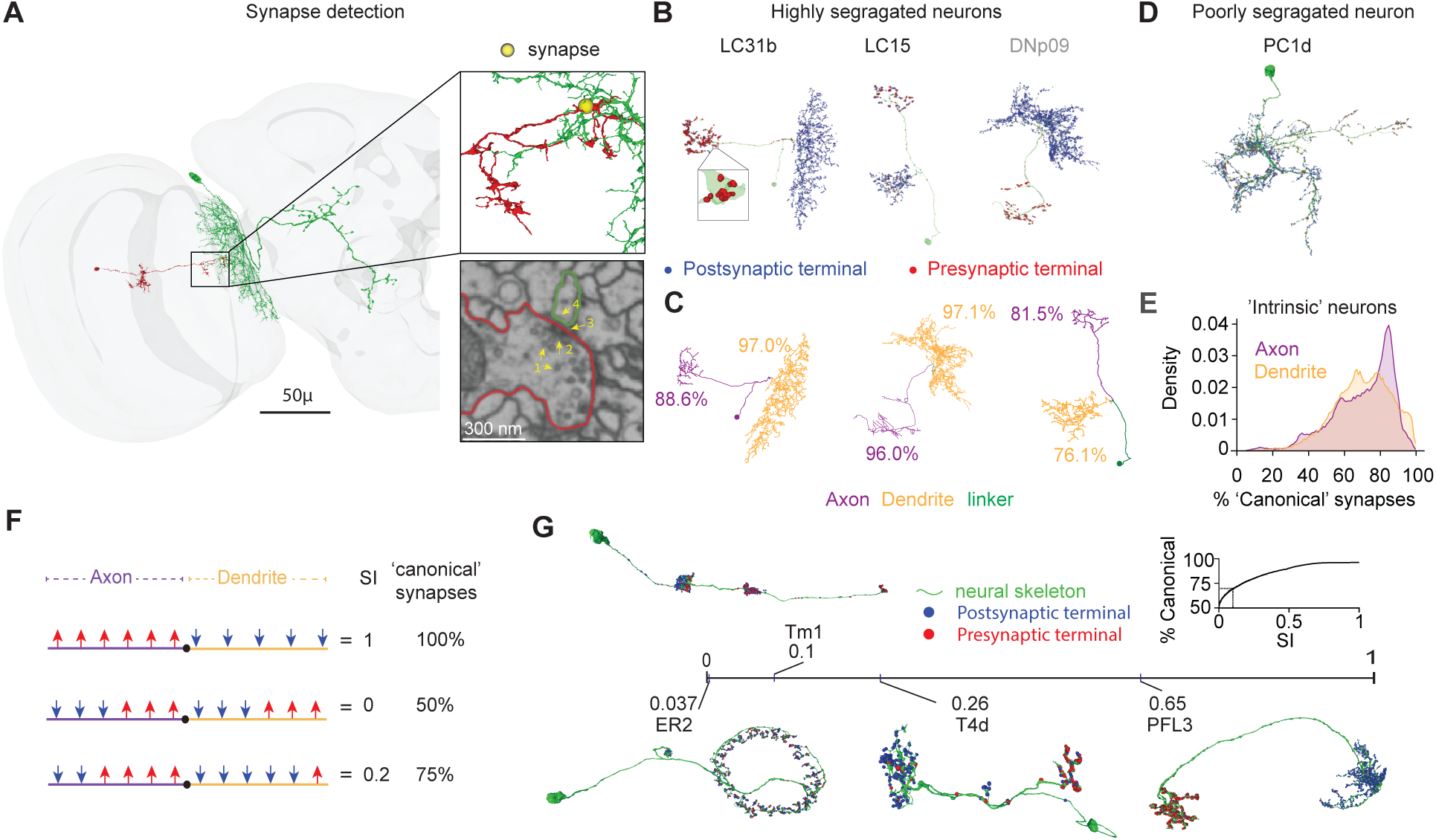
Mixed polarity in dendrites and axons. **A** Two-dimensional view of two connected neurons, and a cross-sectional view of one chemical synapse obtained using Neuroglancer. Visible features are indicated with yellow arrows: presynaptic vesicles (1), presynaptic T-bars (2), synaptic clefts (3), and postsynaptic densities (4). We used a new synapse detector (“Princeton synapses”)^53^ to identify synapse coordinates. **B** Examples of three well-segregated neurons and their presynaptic (red) and postsynaptic (blue) terminals: the visual projection neurons LC15 and LC31b, and the projection neuron DNp09. DNp09 is colored gray as it is not intrinsic to the brain (see ^42^). While descending neurons primarily project outside the brain, the part that remains in the brain is often still well segregated (Figs. S2 and S3). Cell types were obtained from Codex. The LC31b inset highlights a single multisynapse bouton (MSB). **C** Example neurons (the same cells as shown in panel B) after applying the synapse flow centrality (SFC) procedure to split each skeleton into an axonal region (purple), a dendritic region (orange), and (in some cases) a “linker” (green; typically an area with no or very few synapses). The percentage of “canonical synapses” (presynaptic terminals on the axon or postsynaptic terminals on the dendrite) is shown for each compartment. LC31b: dendrites 97.0% (3488/3595), axons 88.6% (1757/1983); LC15: dendrites 76.1% (636/836), axons 81.5% (463/568); DNp09: dendrites 97.1% (5702/5872), axons 96.0% (2487/2591). **D** An example of a poorly segregated neuron (cell type PC1d): pre- and post-synaptic terminals are highly overlapping across the cell. Presynaptic terminals are shown in red, postsynaptic terminals in blue. **E** The fraction of canonical synapses is calculated separately for the dendrites and axons of the intrinsic neurons in the dataset (n = 117,570). The distribution of canonical synapses is shown across all axons (purple) and dendrites (orange) of the intrinsic neuron. **F** Schematics of calculating the segregation index (SI). SI is calculated for a given neuron using entropy, as previously described^40^, based on the distribution of presynaptic (red) and postsynaptic (blue) terminals on the dendrite (orange) and axon (purple). SI ranges between 0 (not segregated, similar ratio between presynaptic and postsynaptic terminals on the dendrite and axon) and 1 (fully segregated, 100% presynaptic on the axon and 100% postsynaptic on the dendrite). Importantly, SI is not a linear function of the overall fraction of canonical synaptic terminals. For example, with 4/6 presynaptic terminals on the axon and 5/6 postsynaptic terminals on the dendrite (9/12 = 75% canonical synapses), SI = 0.2. **G** Examples of individual neurons and their SI values. Green indicates the neuronal skeleton, red indicates presynaptic sites, and blue indicates postsynaptic sites. Inset: percentage of canonical synapses over all the neurons in the dataset as a function of the SI threshold (smoothed with a running average of n = 400 neurons). SI = 0.1 was used as a threshold classifying neurons as non-segregated/not-polarized (SI < 0.1) or segregated/polarized (SI ≥ 0.1), as explained below.

To systematically separate dendritic and axonal compartments, we adapted a previously described algorithm based on synapse flow centrality (SFC; see Methods^40,41^). In brief, for every pair of presynaptic and postsynaptic terminals, paths are traced along the neuronal skeleton, and each node is assigned the number of such paths that pass through it (Supplementary Fig. 1 - S1A). This measure is a synapse-specific version of betweenness centrality^55^, a standard metric in network analysis. Applying SFC partitions each neuron into dendritic and axonal domains and, in some cases, an intermediate region with very few or no synapses, which we term a “linker” (Fig. 1C, Supplementary Fig. 1 - S1). Synapses on linkers comprise less than 1% of all synapses and are excluded from most analyses. In strongly polarized neurons, the vast majority of postsynaptic terminals localize to dendrites and presynaptic terminals to axons (Fig. 1B). Similar results are obtained using an independent synapse detector^54^ (Supp Fig. 1 - S1B-C).

As an initial quantification of mixed polarity across the FlyWire/FAFB connectome, we computed for each neuron the fraction of canonical synapses, defined as postsynaptic terminals on dendrites and presynaptic terminals on axons (Fig. 1E). To avoid biases introduced by neurons that are not fully contained within the brain volume, such as ascending and descending projections to the ventral nerve cord, we focused here and in most subsequent analyses on intrinsic brain neurons (see Table 1). This group includes the central, optic, visual projection, and visual centrifugal superclasses, comprising 118,347 of the 139,255 neurons in FlyWire/FAFB^42^. Across intrinsic neurons, an average of 70.0% of dendritic synapses and 68.9% of axonal synapses are canonical, implying that approximately 30% of all synapses in the adult fly brain are non-canonical. Importantly, the use of SFC assumes that neurons can be meaningfully divided into dendritic and axonal compartments. This assumption breaks down for neurons with highly intermingled synaptic distributions. We address this limitation explicitly in the following section.

Examining canonical synapse fractions across neuronal superclasses reveals substantial heterogeneity both within and between groups (Supplementary Fig. 1 - S1D). Visual centrifugal neurons are among the most strongly segregated neuronal classes. In central and visual projection neurons, both of which project primarily within the brain, dendrites are, on average, more canonical than axons, whereas the opposite trend is observed in optic neurons. These differences are accompanied by systematic differences in the relative prevalence of dendrodendritic and axo-axonic connectivity, as discussed below. Considerable heterogeneity is also present among non-intrinsic superclasses (Supplementary Fig. 1 - S1D), although compartment assignment in incomplete neurons is often unreliable. The correlation between dendritic and axonal canonical fractions varies between superclasses, with the weakest correlation observed in optic neurons (Supplementary Fig. 1 - S1E).

Together, these analyses reveal that mixed polarity is widespread throughout the adult *Drosophila* brain and accounts for a substantial fraction of all synapses. In the following sections, we examine how mixed polarity shapes connectivity patterns across the brain, contributes to reciprocal interactions and local computation, and is implemented by specific subcellular motifs.

### Neuronal polarity varies systematically across developmental stages, cell types, neurotransmitters, and brain regions

To quantify the overall segregation of synapses within individual neurons, we used the entropy-based measure SI^40^ (see Methods). In brief, the SI captures the extent to which presynaptic and postsynaptic terminals are segregated between dendritic and axonal compartments, with SI = 1 corresponding to a fully segregated neuron and SI = 0 indicating identical pre-/post-synaptic ratios in dendrites and axons (Fig. 1F). SI values were robust to the choice of synapse detector and were highly reproducible between left–right homologs and among neurons belonging to the same cell type (Supplementary Fig. 2–S1–S2; see Methods). We also tested whether excluding synapses onto small unproofread segments (“twigs”) biased SI estimates. Including these synapses had only a minor effect on SI values (Supplementary Fig. 2–S1A). Additionally, we found no significant correlation across neuropils between the fraction of synapses connected to unproofread segments and the fraction of AD, AA, DD, or DA synapses (data not shown), suggesting that exclusion of unproofread segments does not substantially bias synapse-type estimates.

Illustrative examples span a broad range of polarity. Central complex ring neurons (ER2) exhibit very low SI values, PFL3 neurons are highly segregated, and optic neurons such as Tm1 and T4d show intermediate segregation (Figure 1G). Importantly, SI is not a linear measure of canonical connectivity: neurons with SI = 0.1 still contain approximately 70% canonical synapses (Figure 1F, inset).

Across all intrinsic neurons in FlyWire/FAFB, the SI distribution is bimodal, with a large fraction of neurons exhibiting extensive mixed polarity (46% of neurons have SI < 0.1; Fig. 2A). To test whether mixed polarity in flies is primarily a consequence of small nervous system size relative to vertebrates, we examined the SI distribution in the larval L1 connectome^41^. Although mixed-polarity neurons are also present in the larvae, L1 neurons are substantially more polarized than adult neurons (Fig. 2A). Mixed polarity is therefore not simply a consequence of nervous system size, but instead reflects more complex developmental and/or functional factors.

**Fig. 2:**
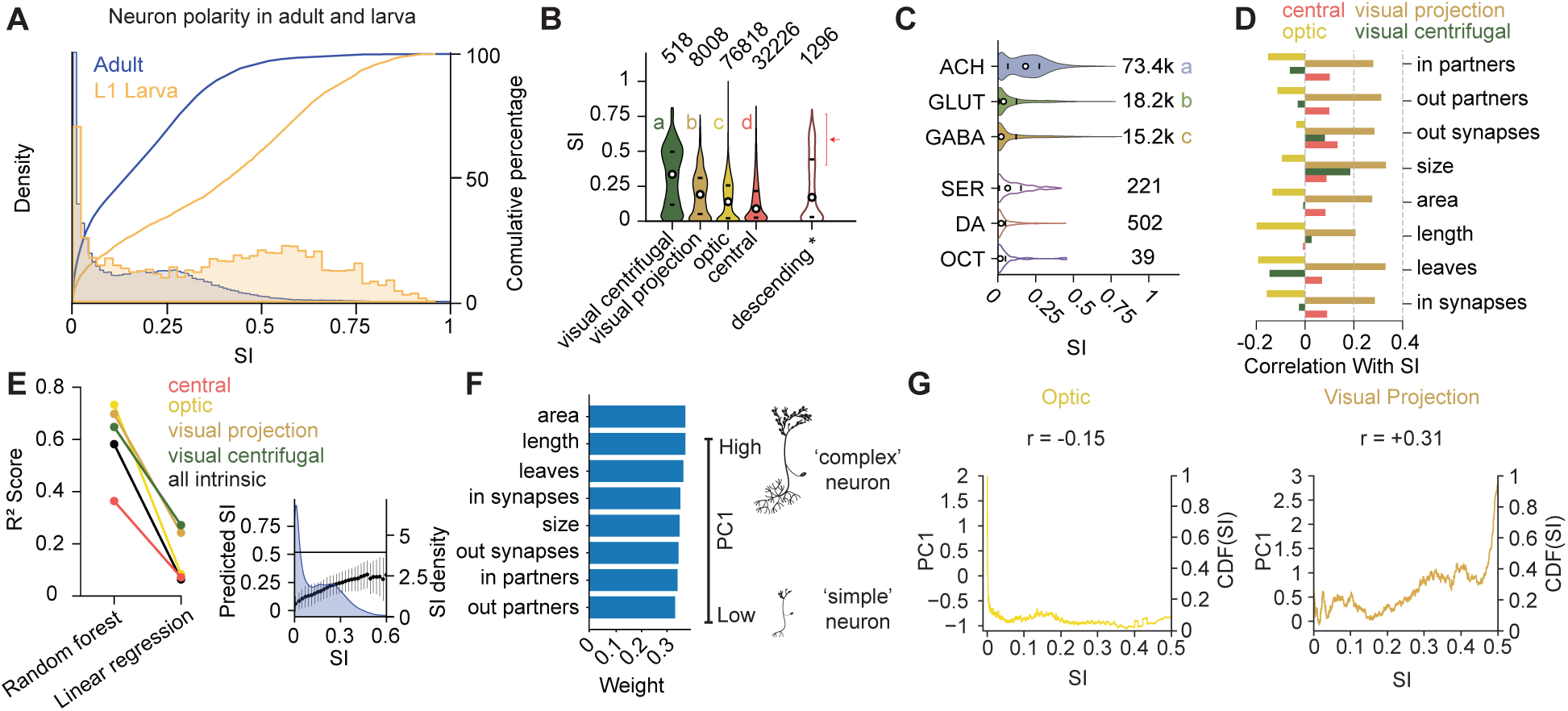
Mixed polarity across the fly brain. **A** SI distribution (left y-axis) and cumulative percentage (right y-axis) for the L1 larva (orange; n = 2,650 neurons) and the adult brain (purple; n = 117,570 intrinsic neurons). For example, while approximately half of larval neurons have SI > 0.4, only 5.7% of adult intrinsic neurons do. **B** Violin plots showing SI distributions across the four intrinsic neuron superclasses. Dots indicate medians and vertical lines show the interquartile ranges; population size is indicated above each violin. Group differences across super-classes were significant (Kruskal–Wallis, p < 0.001), with all Dunn’s post-hoc pairwise comparisons remaining significant after multiple-comparison correction). The red arrow indicates high SI values for descending neurons, despite most of their outputs being absent from the dataset (see examples in Fig. 1B and Supp Fig. 2 - S3B). **C** Violin plots showing segregation index (SI) distributions across neurotransmitter types for intrinsic neurons in the dataset that have a predicted neurotransmitter (n = 104,174). ACH, acetylcholine; GABA, gamma-aminobutyric acid; GLUT, glutamate; SER, serotonin; DA, dopamine; OCT, octopamine. Points indicate medians and horizontal lines indicate the interquartile ranges. Among the three major neurotransmitter classes analyzed statistically (ACH, GABA, and GLUT), SI differed significantly between groups (Kruskal–Wallis, H(2) = 15118.97, p < 0.001, η² = 0.141). Bonferroni-corrected Dunn’s post-hoc comparisons were significant for all pairwise contrasts. Effect sizes indicated larger differences for ACH–GABA and ACH–GLUT, and a smaller effect for GABA–GLUT (see Table 1). **D** Correlations between SI and selected neuronal features across intrinsic superclasses. **E** Model performance for predicting SI using the eight indicated features. The coefficients of determination (R²) for linear regression and random forest models are shown, computed separately for each superclass and for all superclasses combined (black). Right: SI predicted by the random forest model plotted against observed SI (binned). The points denote the mean predicted SI within each SI bin, and the vertical bars indicate the standard deviation of predictions across trees in the random forest. The blue curve shows the overall SI distribution for intrinsic neurons. The model underestimates the SI value for large SI values (which are rare in the dataset). **F** Weights of the eight features in the first principal component (PC1) derived from principal component analysis (PCA) of standardized neuronal features (see Methods). Bars represent the signed weight of each feature in PC1; larger positive values indicate stronger positive contributions. PC1 explained 87% of the total variance, indicating that the eight features largely capture a single axis ranging from small, weakly connected neurons to large, highly connected neurons. Inset: because all features contribute positively to PC1, higher PC1 values correspond to larger, more connected neurons (“complex”), whereas lower PC1 values correspond to smaller, less connected neurons (“simple”). **G** The correlation between SI and PC1 for optic neurons is negative (top, r = −0.15), while for visual projection neurons it is positive (bottom, r = +0.31). Colored lines show the rolling mean (window size = 400) of PC1 as a function of SI.

We next examined how segregation varies across neuronal superclasses. Consistent with differences in canonical synapse fractions (Supp Fig. 1 - S1D), neurons linking the optic lobes and central brain are, on average, more polarized than neurons whose arbors are largely confined to either the central brain or optic lobes (Fig. 2B). This increased polarity may reflect feedforward information flow between distant brain regions. In contrast, the SI distributions of most non-intrinsic superclasses are biased toward low values, likely because major input or output arbors lie outside the reconstructed FlyWire/FAFB volume (Supp Fig. 2–S3)^42^.

Interestingly, despite lacking their primary ventral nerve cord (VNC) projections, descending neurons exhibit relatively high SI values (mean SI = 0.20) compared with central neurons (mean SI = 0.163). This can be explained by the fact that many descending neurons possess substantial output arbors within the brain, often in the subesophageal zone (SEZ; see examples in Supplementary Fig. 2–S3B), consistent with recent evidence that descending neurons form dense local networks within the brain^56^.

We next asked whether neuronal polarity differs across neurotransmitter classes. Cholinergic neurons, which are predominantly excitatory, are significantly more segregated than either GABAergic or glutamatergic neurons (Fig. 2C). The higher polarity of cholinergic neurons may reflect feedforward connections linking distinct neuropils, in contrast to the more local computations often mediated by inhibitory neurons. An extreme example of mixed polarity and local computation is provided by the GABAergic CT1 optic neuron^57^.

We then asked whether neuronal polarity is related to intrinsic cellular properties, including morphology and connectivity. We quantified eight neuronal features: four morphological (size, area, length^42^, and number of leaves) and four connectivity-related (number of input partners, output partners, input synapses, and output synapses). Many of these features were strongly correlated with one another (Supplementary Fig. 2–S4A), suggesting that they largely capture a common axis of neuronal complexity. Nevertheless, their relationships with SI differed across neuronal superclasses (Fig. 2D). In central and visual projection neurons, greater segregation was generally associated with larger and more connected neurons. In contrast, the opposite trend was observed in optic neurons, where small, highly polarized neurons were common.

Although SI exhibited significant linear correlations with multiple cellular features, prediction of SI was substantially improved using a random forest regression model (Fig. 2E), indicating that the relationship between neuronal properties and polarity is partly nonlinear. To summarize the eight features in a single variable, we applied principal component analysis (PCA). The first principal component (PC1) explained 87% of the total variance and assigned positive weights to all eight features (Fig. 2F), indicating that it captures a continuum from small, sparsely connected neurons to large, highly connected neurons. We therefore refer to PC1 as neuronal complexity.

The relationship between complexity and polarity differed across neuronal superclasses. For central and visual projection neurons, higher complexity was associated with increased segregation. For optic neurons, we observed an overall negative relationship between complexity and SI, largely due to the existence of complex neurons (large PC1) with very low SI (Fig. 2G; Supplementary Fig. 2–S4C–D). For example, MeTu1 visual projection neurons are small and weakly segregated (mean SI = 0.00014), whereas the larger and more complex LC11 neurons are strongly polarized (mean SI = 0.35), potentially reflecting the need for high polarity in neurons that convey information from the optic lobe to distant brain regions. In contrast, within the optic lobe, T4c neurons are relatively small yet highly polarized, whereas the much larger Li26 neurons exhibit strong mixed polarity (Supplementary Fig. 2–S4C). This pattern may reflect the highly ordered, layered organization of the optic lobe, which contains many small but strongly polarized neurons within individual visual processing layers^58^.

Taken together, these analyses show that mixed polarity varies systematically with developmental stage, neuronal superclass, neurotransmitter identity, and cellular complexity. The strong dependence of polarity on neuronal identity and organization suggests that mixed polarity is unlikely to reflect developmental or wiring noise but rather reflects functionally meaningful circuit architectures.

### Mixed polarity underlies widespread non-canonical connectivity

Mixed polarity provides the structural basis for non-canonical connectivity AA synapses require postsynaptic sites on axons, and DD synapses require presynaptic sites on dendrites. We next characterized how neurons in the fly brain are connected via canonical AD synapses and non-canonical AA, DD, and DA connections.

Highly mixed cells, such as many of the “ring” neurons in the central complex and the “Dm” neurons of the *Drosophila* optic medulla that may function as multi-output devices^59^ (Fig. 3A), lack clearly separable axonal and dendritic arbors. Therefore, to reliably assign synapses to axons or dendrites, we restricted subsequent analyses to polarized neurons, defined as cells with SI ≥ 0.1 (Fig. 3A). While this threshold is somewhat arbitrary, it approximately splits the bimodal distribution of SI into non-polarized neurons (peak close to SI = 0) and polarized neurons (peak around 0.27; Fig. 2A). In polarized neurons, four types of synaptic connections can occur (Fig. 3B): canonical AD connections and three non-canonical types: AA, DD, and DA. Using SI = 0.1, we defined a “mixed compartment” as a dendrite with <67.3% postsynaptic terminals or an axon with <71% presynaptic terminals (Supplementary Fig. 3 - S1A).

**Fig. 3:**
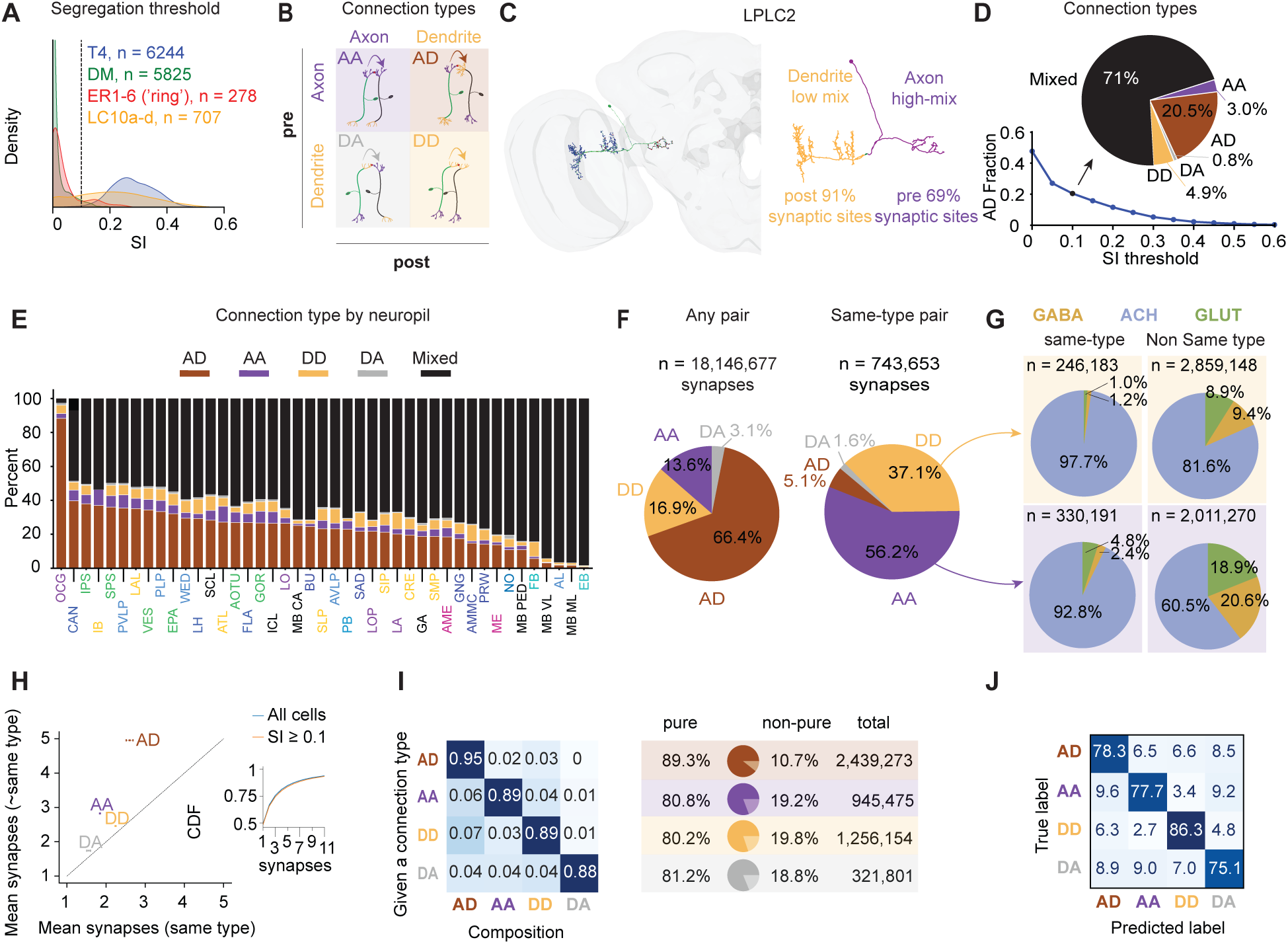
Connection types across FlyWire. **A** SI distributions for four cell types: T4, Dm, Ring, and LC10a-d lobula columnar neurons. Vertical dashed line: SI = 0.1. **B** Schematics of the four possible connection types: AD, AA, DD, and DA. Each configuration is illustrated with a presynaptic neuron (green) and a postsynaptic neuron (black). **C** An example neuron (cell type LPLC2) with a non-mixed dendrite (>67.3% postsynaptic terminals; see Supplementary Fig. 3–S1) and a mixed axon (<71% presynaptic terminals), using an SI threshold of 0.1. **D** Bottom: fraction of canonical (AD; axo-dendritic) synapses as a function of the SI threshold. Increasing the threshold imposes stricter canonicality criteria, thereby increasing the fraction of compartments classified as mixed and, consequently, the fraction of synapses classified as mixed. At SI = 0, no compartments are labeled mixed. Using a threshold of SI = 0.1, 71% of all synapses are classified as mixed, meaning that their presynaptic site, postsynaptic site, or both are located in mixed compartments. In contrast, 20.5% of all synapses are classified as AD, occurring between a non-mixed axon and a non-mixed dendrite. Top: fraction of synapses in each category using an SI threshold of 0.1. **E** Distribution of synapse types across neuropils (n = 65,952,300 synapses). **F** Proportions of synapse types connecting pairs of polarized neurons (SI ≥ 0.1) for all pairs (n = 18,146,677 synapses; left) and for same-type pairs (n = 743,653 synapses; right). **G** Predicted neurotransmitters (NTs) of presynaptic neurons in axo-axonic (top) and dendrodendritic (bottom) connections between neurons of the same type (left) and between neurons of different types (right). **H** Comparison of mean connection strength (number of directed synapses) involving same-type versus different-type pairs; inset: cumulative distribution of synapse counts for all neurons and for polarized neurons (SI ≥ 0.1), indicating that approximately 80% of connected pairs have five synapses or fewer. The points represent means and the error bars indicate SEM along both axes. Error bars may not be visible if they are smaller than the symbol size. **I** Left: conditional probabilities of synaptic-type composition. For example, among all connections containing at least one AD synapse (top row), 95% of synapses are AD, 2% are AA, and 3% are DD. Right: fraction of “pure” connections for each connection type, given the presence of at least one synapse of that type. For example, among all directed neuron pairs containing at least one AD synapse, 89.3% contain only AD synapses. Connections containing multiple synapse types (e.g., both AA and DD; see examples in Fig. 6E, F) are not considered pure. **J** Confusion matrix for the random forest model predicting synapse type using the eight features of both the presynaptic and postsynaptic neurons (16 features in total).

Notably, many neurons have one non-mixed and one highly mixed compartment (see e.g., the visual projection LPLC1 neurons; Fig. 3C). Under these criteria, only 20.5% of all synapses correspond to canonical AD connections between a non-mixed axon and a non-mixed dendrite (Fig. 3D). This result is robust to the choice of synapse detector (Buhmann; Supplementary Fig. 3 - S1A). Varying the SI threshold predictably shifts this estimate: at SI = 0.05, 27.1% of synapses are classified as canonical; this decreases to 11.6% at SI = 0.2 (Fig. 3D). An SI of around 0.05 often includes clearly non-polarized neurons (Fig. 3A), and SI ≥ 0.2 excludes a large fraction of otherwise well-segregated cells (only 35.4% of intrinsic neurons meet this criterion).

We find that the prevalence of synapse types varies substantially across neuropils (Fig. 3E; similar results are obtained using the Buhman synapse detector, Supplementary Fig. 3 - S2A-C). Several regions, including the central complex (comprising the ellipsoid body [EB], fan-shaped body [FB], and noduli [NO]), antennal lobe (AL), and medial and ventral lobes of the mushroom body (MB ML, MB VL), are particularly enriched in non-canonical synapses, consistent with prior connectomic studies describing recurrent motifs in the central complex, DD connectivity in olfactory circuits, and AA interactions among mushroom body output pathways^60–62^. From this point onward, all analyses of connection types (AD, AA, DD, and DA) include only polarized neurons (SI ≥ 0.1), for which axonal and dendritic compartments are generally well defined.

Across polarized neurons, 33.6% of all connections are non-canonical (AA, DD, or DA; Fig. 3F, left). This fraction increases dramatically for connections between neurons of the same cell type (Fig. 3F, right): more than 93% of synapses between same-type neurons are AA or DD, consistent with shared axonal or dendritic domains. The fraction of cholinergic connections is larger for AA and DD connections between same-type pairs compared with pairs belonging to different cell types, possibly reflecting a bias toward lateral excitation between cells that share similar functions. On average, same-type connections are slightly weaker than connections between different types (Fig. 3H).

Connections between a given presynaptic neuron A and postsynaptic neuron B can involve multiple synapse types (Fig. 3I), although one type typically dominates. For example, among all neuron pairs with at least one DD synapse, just 7% of the synapses are AD (Fig. 3J), and in 13% of such pairs, there is at least one AA synapse (Supplementary Fig. 3 - S2D). Overall, only around 80% of non-canonical synapses (AA, DD, and DA) are “pure” (i.e., involve a single connection type), compared with approximately 90% of canonical AD connections. Reciprocal connectivity, which may involve the same or different connection types in opposite directions, is not analyzed here and is addressed in the next section.

Last, we asked whether synapse type can be predicted from the morphology and connectivity properties of pre- and post-synaptic neurons. Using eight properties (as in Fig. 2F), we were able to predict synapse type with >75% accuracy, compared with a chance level of 25% (Fig. 3J; see Table 4). Morphology and connectivity-related features of pre- and post-synaptic neurons contributed equally to the classification (Supplementary Fig. 3 - S2E).

Taken together, our analysis reveals that non-canonical connectivity is widespread across the adult *Drosophila* brain. Synapse type varies by brain region, and by the properties and the similarities between the pre- and post-synaptic partners. In the next section, we examine how the properties of the pre- and post-synaptic cells predict the connection type.

### A simple architectural rule predicts synapse type

We next asked whether properties of presynaptic and postsynaptic neurons could predict synapse type (AD, AA, DD, or DA). To do so, we used the first principal component (PC1) introduced above, which summarizes eight morphological and connectivity-related features and explains 87% of their variance (Fig. 2F). As discussed above, we interpret PC1 as a measure of neuronal complexity.

Remarkably, a model using only the PC1 values of the presynaptic and postsynaptic neurons predicts synapse type with relatively high accuracy (Fig. 4A). Although performance is lower than that of a model using all eight features for both neurons (Fig. 3J), the reduction in accuracy is only approximately 10%. Thus, much of the information required to predict synapse type is captured by a single measure of neuronal complexity. Prediction accuracy remains lower than that achieved when the identities of the presynaptic and postsynaptic cell types are known (Fig. 4B), indicating that cell-type-specific information also contributes to determining synapse type.

**Fig. 4:**
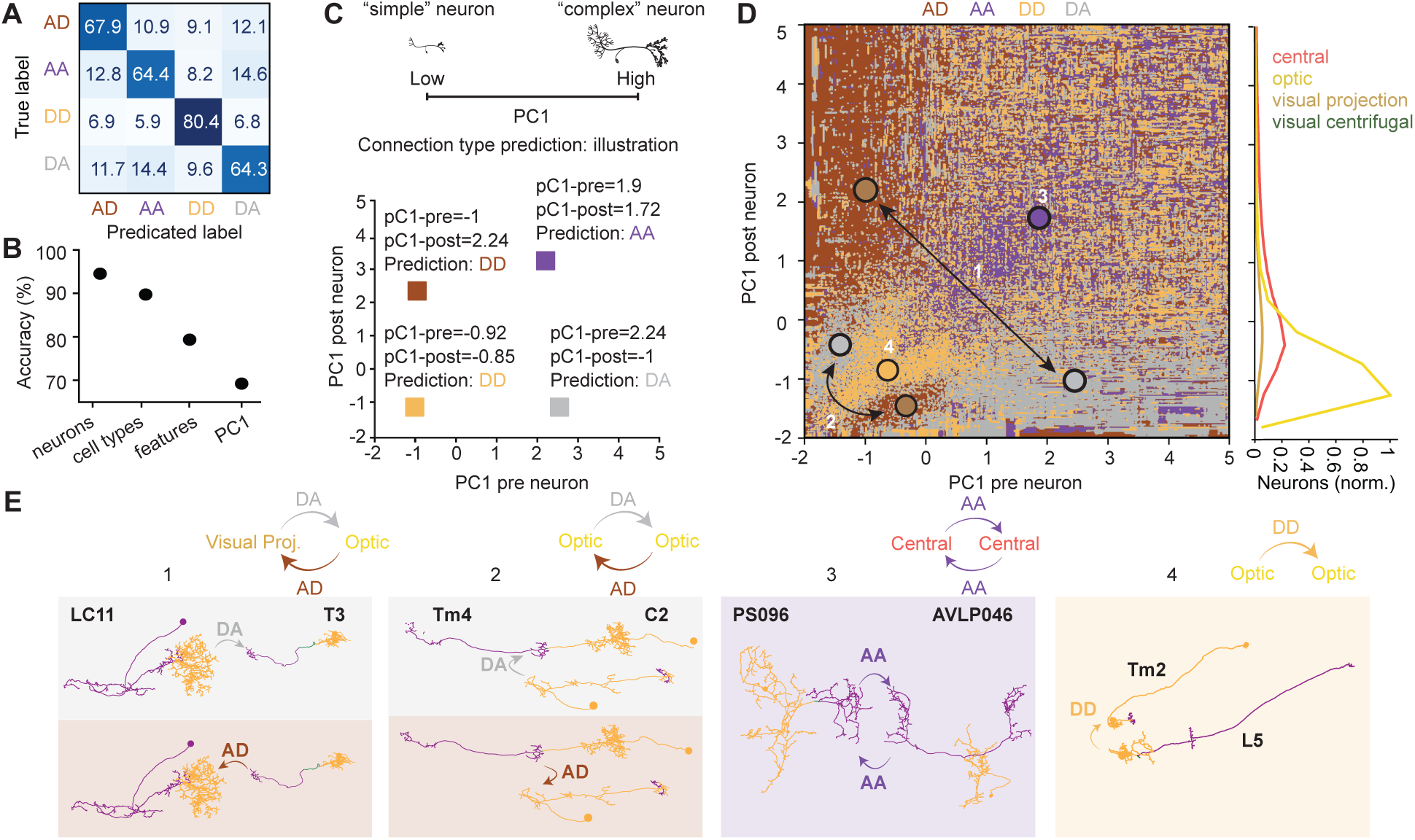
Synapse type prediction based on pre- and post-synaptic partners. **A** Confusion matrix for a model trained using only the first principal component (PC1) of the pre- and post-synaptic neurons. **B** Accuracy of predicting synapse type (for each individual synapse) using four different models (see Methods): (1) prediction based on the identity of the presynaptic and postsynaptic cells. The type of each synapse (AD, AA, DD, and DA) is predicted as the most common synapse type observed for that specific neuron pair. (2) Prediction based on the cell types of the pre- and post-synaptic neurons. The model assigns each synapse the most frequent synapse type observed among all pairs belonging to those presynaptic and postsynaptic cell types, (3) a feature-based classifier (see Fig. 3J), and (4) a reduced model using only two features: PC1 of the pre- and post-synaptic cells (panel A). Model (4) uses only PC1 of the presynaptic and postsynaptic neurons, which together summarize the eight neuronal features shown in Fig. 2F. The confusion matrix in Fig. 4A and the decision surface in Fig. 4D are both for model (4). **C** Illustration of the decision surface shown in D. For a synapse between a presynaptic neuron with PC1 = 1.9 and a postsynaptic neuron with PC1 = 1.72, the model predicts a connection type AA. For reciprocally connected neurons, the presynaptic and postsynaptic PC1 values are exchanged between the two directions of the connection. In the illustrated example (brown and gray boxes), a pair of neurons with PC1 values of −1 and 2.24 is reciprocally connected. For the connection from the neuron with PC1 = −1 to the neuron with PC1 = 2.24, the model predicts an AD synapse. Reversing the direction of the connection exchanges the two PC1 values (2.24, −1), and the model predicts a DA synapse. Thus, exchanging the presynaptic and postsynaptic neurons (equivalently, reflecting a point across the diagonal y = x) exchanges AD and DA predictions, as observed in panel D. **D** Decision surface of the model predicting synapse type from the PC1 values of the presynaptic and postsynaptic neurons (see illustration in C). Numbered circles indicate the examples shown in E: (1) an AD–DA reciprocal connection between a visual projection neuron (LC11, PC1 = 2.24) and an optic neuron (T3, PC1 = −1); (2) an AD–DA reciprocal connection between two optic neurons (Tm4 and C2; PC1 = −0.43 and −1.39, respectively); (3) a reciprocal AA connection between two central neurons (AVLP046 and PS096; PC1 = 1.9 and 1.72, respectively); and (4) a DD connection between two optic neurons (L5 and Tm2; PC1 = −0.92 and −0.85, respectively). Right panel: distribution of PC1 values across neurons in each superclass. **E** Skeletons of the neurons shown in D, colored by compartment: axons in purple and dendrites in orange. The corresponding synapse type is illustrated for each example. Note that the example neurons PS096 and AVLP046 are reciprocally connected through AA synapses.

Before examining the full decision surface, it is useful to consider how reciprocal connections are represented in this framework (Fig. 4C). The model predicts synapse type from two values: the complexity of the presynaptic neuron (PC1-pre) and the complexity of the postsynaptic neuron (PC1-post). Exchanging the direction of a connection simply swaps these two values. Thus, for a reciprocally connected pair of neurons, the two directions of the connection correspond to two points that are mirror images of one another across the diagonal (y = x) of the decision surface. For example, a connection from a simple neuron (PC1 = −1) to a complex neuron (PC1 = 2.24) is predicted to be AD, whereas the reciprocal connection is predicted to be DA. Consequently, AD and DA occupy mirror-image regions of the decision surface. As shown in the next section, reciprocal AD↔DA motifs are among the most common forms of reciprocal connectivity in the fly brain.

The resulting decision surface reveals a striking organization of synapse types in the space defined by presynaptic and postsynaptic complexity (Fig. 4D). Two broad regimes emerge. The first regime is dominated by simple neurons (PC1 < 0.5), which are primarily optic neurons, the largest neuronal superclass in the fly brain. In this regime, neurons of similar complexity tend to connect through DD synapses (for example, the L5–Tm2 pair; Fig. 4D, E). Connections between neurons of different complexity are predicted to be AD or DA, depending on the direction of the connection (for example, the Tm4–C2 pair; Fig. 4D, E). Interestingly, this organization is not restricted to visual circuits. The antennal lobe, an early olfactory processing center, and the antennal mechanosensory and motor center (AMMC), an early mechanosensory processing center, are also enriched in DD connectivity (see Fig. 5I), suggesting that dendrite-based interactions may represent a general feature of early sensory processing circuits.

**Fig. 5:**
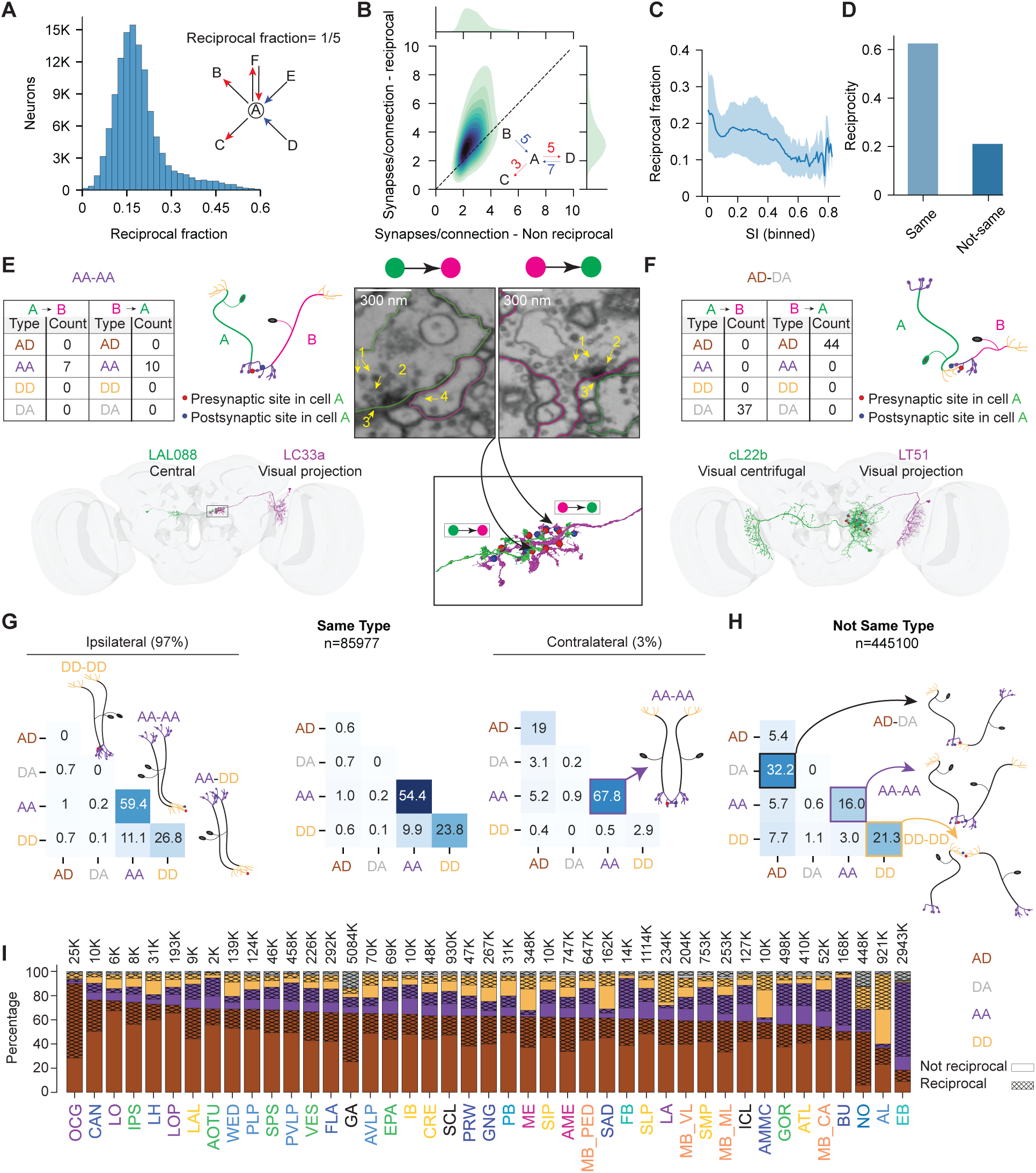
Mixed polarity underlies reciprocal connectivity. **A** Distribution of the reciprocal fraction across intrinsic neurons in FlyWire/FAFB (n = 118,347). The reciprocal fraction is defined as the proportion of a neuron’s partners with which it forms bidirectional connectivity (see inset; an arrow represents an existing connection, with any number of synapses). **B** Reciprocal connections contain more synapses per connection than non-reciprocal connections (n = 117,407 neurons). Inset: neuron A has two non-reciprocal partners (B and C), represented by two directed edges containing a total of eight synapses, yielding a mean of 8/2 = 4 synapses per directed connection. Neuron A also has one reciprocal partner (D), represented by two directed edges containing a total of 12 synapses, yielding a mean of 12/2 = 6 synapses per directed connection. **C** Mean reciprocal fraction as a function of the segregation index (SI), showing higher reciprocity in mixed (less segregated) neurons. **D** Fraction of connected neuron pairs that are reciprocal, grouped according to whether the two neurons belong to the same cell type or different cell types. **E, F** Examples of reciprocal motifs composed of AD↔DA and AA↔AA synaptic configurations (see also the DD↔DD example in Supplementary Fig. 5–S1). Each table shows the synaptic composition in both directions of a reciprocal pair. For the AA↔AA example, a synaptic cross-section is shown for one synapse in each direction. In the AD↔DA example, the ratio of AD to DA synapses is 44:37. Across all AD↔DA reciprocal motifs in the dataset, the mean ratio was 4.4:1. **G** Dominant synaptic-type compositions of reciprocal connections (>50% of synapses in each direction) for neuron pairs belonging to the same cell types, separated into ipsilateral (97% of pairs) and contralateral (3% of pairs) connections. **H** Dominant synaptic-type compositions of reciprocal connections for neuron pairs belonging to different cell types. For example, 32.2% of reciprocally connected non-same-type pairs are connected through an AD connection in one direction and a DA connection in the other. **I** Proportions of synapses in each neuropil belonging to the different connection types. Reciprocal connections are indicated by diamond-mesh shading. For example, the antennal lobe and the antennal mechanosensory and motor center, early olfactory and mechanosensory processing regions, respectively, are enriched in DD connections, approximately half of which are reciprocal. In contrast, the mushroom body calyx, the main olfactory input region of the mushroom body, is enriched in AA connections, approximately half of which are reciprocal. Across neuropils, the majority of DA synapses form part of reciprocal connections.

A different pattern emerges in the second regime, where at least one of the two neurons is relatively complex (PC1 > 0.5). These neurons are predominantly central neurons. Here, neurons of similar complexity tend to connect through AA synapses (for example, the AVLP046–PS096 pair; Fig. 4D, E). As in the first regime, connections between neurons of different complexity are predicted to be AD or DA in opposite directions (for example, the LC11–T3 pair; Fig. 4D, E). Thus, moving along the diagonal of the decision surface reveals a transition from DD-dominated interactions between simple neurons to AA-dominated interactions between complex neurons.

The prevalence of DD and AA synapses along the diagonal of the decision surface is consistent with our earlier observation that same-type neurons are predominantly connected through DD and AA synapses (Fig. 3F).

Taken together, these results suggest a simple first-order architectural principle. Small neurons, which are enriched in early sensory circuits across multiple modalities, tend to interact through their input compartments (dendrites), whereas larger downstream neurons, enriched in central brain regions, tend to interact through their output compartments (axons). Thus, a single axis of neuronal complexity is sufficient to explain a substantial fraction of the organization of synapse types across the fly brain.

### Reciprocal connectivity emerges from non-canonical synaptic motifs

Reciprocal connections, i.e., pairs of neurons connected in both directions, are a well-established feature of brain networks and are consistently overrepresented relative to random graph models such as Erdős–Rényi and configuration networks^45–47^. Comparative studies further suggest that the overall level of reciprocity in the *Drosophila* brain is comparable with that observed in other nervous systems, including the complete connectomes of hermaphrodite and male *C*. *elegans*^22,48,49^, as well as subvolume reconstructions of the larval zebrafish hindbrain^50^ and mouse visual cortex^51,52^.

To examine how mixed polarity contributes to reciprocity, we defined the “reciprocal fraction” of a neuron as the fraction of its synaptic partners with which it forms bidirectional connectivity (Fig. 5A). Across the FlyWire/FAFB connectome, an average of 19.2% of a neuron’s partners are reciprocal. Reciprocal connections are also stronger than non-reciprocal connections, on average containing more synapses per directed connection (Fig. 5B).

Reciprocal connectivity is heavily dependent on neuronal polarity. Highly mixed neurons (SI ≈ 0) have approximately twice the reciprocal fraction of highly polarized neurons (Fig. 5C), indicating that mixed polarity strongly promotes bidirectional interactions. In addition, connected neuron pairs belonging to the same cell type are approximately three times more likely to be reciprocal than pairs belonging to different cell types (Fig. 5D).

We next asked how different synapse types contribute to reciprocal connectivity. Reciprocal interactions can be mediated by one or more combinations of canonical and non-canonical synapse types. For example, a pair of neurons of types LAL088 (central) and LC33a (visual projection) is reciprocally connected exclusively through AA synapses, with seven synapses in one direction and ten in the other (Fig. 5E). As in this example, reciprocal synapses are often located in close spatial proximity. Similarly, reciprocal interactions frequently occur between dendrites (DD↔DD; Supplementary Fig. 5–S1A). In another example, two LC11 neurons are reciprocally connected through both their axons and dendrites, simultaneously forming AA↔AA and DD↔DD motifs (Supplementary Fig. 5–S1B). In yet another example, the axon of the projection neuron cL22b and the dendrite of the visual centrifugal neuron LT51 are reciprocally connected through complementary AD and DA synapses (Fig. 5F). Across all AD↔DA reciprocal motifs, canonical AD synapses outnumber non-canonical DA synapses by an average ratio of 4.4:1. This imbalance suggests that many AD↔DA motifs correspond to predominantly feedforward pathways supplemented by local reciprocal feedback. Manual validation of synapse direction indicated that approximately 80% of neuron pairs containing one or more detected DA synapses are expected to contain at least one true DA synapse, supporting the biological relevance of these motifs (see Methods).

The composition of reciprocal motifs depends strongly on the relationship between the two neurons (Fig. 5G, H). Reciprocal connections between neurons of the same cell type are dominated by AA↔AA motifs, followed by DD↔DD motifs. Approximately 10% of same-type reciprocal pairs are connected asymmetrically, with AA synapses in one direction and DD synapses in the other. These motifs often arise when neurons overlap extensively within their axonal or dendritic domains (Supplementary Fig. 5–S1B). A small fraction (approximately 3%) of same-type reciprocal pairs are contralateral. Among these mirror-neuron pairs, 67.8% of reciprocal connections are AA↔AA, whereas only 2.9% are DD↔DD, indicating that contralateral reciprocity occurs predominantly within output domains.

A different pattern is observed for reciprocal connections between neurons of different cell types. Here, the most common motif is AD↔DA, followed by AA↔AA and DD↔DD connections (Fig. 5H). Notably, purely canonical reciprocal motifs (AD↔AD) are rare, accounting for just 5.4% of reciprocal connections between different-type neurons and 0.6% between same-type neurons. Thus, most reciprocal connectivity in the fly brain is mediated through non-canonical synaptic motifs.

The prevalence of reciprocal motifs also varies widely across brain regions (Fig. 5I). The antennal lobe and AMMC, early olfactory and mechanosensory processing centers, respectively, are enriched in reciprocal DD↔DD connectivity. In contrast, the mushroom body calyx, a higher-order olfactory region, and the EB, a sensorimotor center involved in navigation, are enriched in reciprocal AA↔AA motifs.

Interestingly, the antennal lobe is often considered the insect analog of the mammalian olfactory bulb, a structure well known for its extensive DD circuitry^19^, suggesting that recurrent DD connectivity may represent a conserved solution for early sensory processing across phyla.

Together with the complexity-based organization described in Fig. 4, these observations suggest a broader architectural principle. At the cellular level, simple neurons (low PC1), which are predominantly optic neurons and therefore enriched in early stages of visual processing, preferentially participate in DD interactions. At the circuit level, early sensory regions such as the AL and AMMC are enriched in reciprocal DD↔DD motifs, whereas higher-order and integrative regions are enriched in AA↔AA motifs. These observations raise the possibility that DD interactions are preferentially associated with early sensory processing, whereas AA interactions become increasingly prominent in downstream and integrative circuits.

Together, these findings show that reciprocity in the fly brain is closely linked to mixed polarity and is mediated predominantly through non-canonical synaptic motifs. The specific form of reciprocity depends on both neuronal identity and brain region, revealing a rich repertoire of reciprocal architectures that extend far beyond canonical AD↔AD connectivity.

### Different circuit classes implement distinct reciprocal motifs

To examine how mixed polarity shapes connectivity within defined neuronal populations, we compared three well-characterized systems: antennal lobe projection neurons (ALPNs), which convey primary olfactory information; lobula columnar (LC) visual projection neurons, which transmit visual information from the optic lobe to the central brain; and central complex (CX) neurons, which participate in higher-order sensorimotor computations.

ALPNs exhibit abundant within-type AA connectivity, consistent with previous observations^27^, and are also rich in DD connections (Fig. 6A; Supplementary Fig. 6–S1A). In contrast, within-type connectivity among visual projection neurons is almost exclusively dominated by AA connections (Fig. 6A; Supplementary Fig. 6–S1B). Because connectivity between same-type optic neurons is predominantly DD (Fig. 5G), this shift toward AA connectivity may reflect the greater distance of visual projection neurons from the sensory periphery compared with optic neurons. Alternatively, it may reflect modality-specific differences between visual and olfactory processing. Within-type connectivity among CX neurons is likewise dominated by AA connections (Fig. 6A; Supplementary Fig. 6–S2), consistent with their role as downstream integrative neurons.

**Fig. 6:**
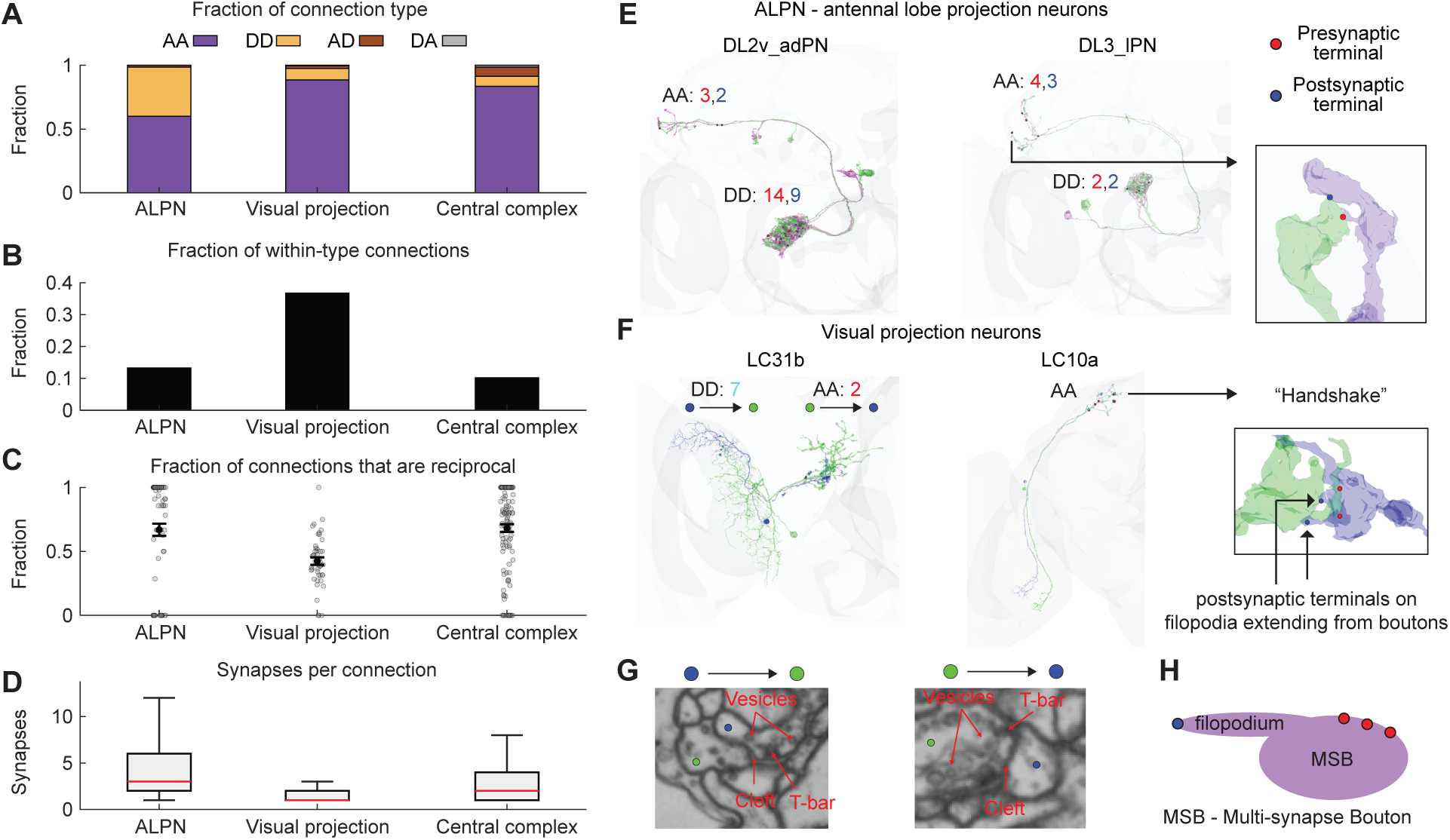
Distinct reciprocal motifs characterize different neural systems. **A** Distribution of connection types. Results are shown separately for antennal lobe projection neurons (ALPNs), visual projection neurons, and central complex neurons. **B** Bar plot showing the fraction of connections that occur between neurons of the same cell type for the same groups shown in A. **C** Fraction of connections that are reciprocal (i.e., a connection is also present in the opposite direction). Each circle represents a single cell type. Means ± SEM are shown. **D** Number of synapses per connection for the same groups shown in A. The central red line indicates the median; the lower and upper edges of the box indicate the 25th and 75th percentiles, respectively. Whiskers extend to the most extreme data points. **E** Example connected neuron pairs. The DL2v–adPN pair exhibits both AA↔AA and DD↔DD reciprocal connectivity. Red and blue circles indicate presynaptic and postsynaptic terminals, respectively, from the perspective of the green neuron. The DL3–IPN pair is likewise reciprocally connected through both axonal and dendritic compartments. Inset: a postsynaptic terminal located on a bouton of the green neuron contributes to local reciprocal connectivity. This motif (a postsynaptic terminal on a bouton) is discussed further below. **F** Same as in E, but for lobula columnar (LC) visual projection neurons. The LC31b pair exhibits an AA↔DD reciprocal motif, with connectivity occurring through dendritic compartments in one direction and axonal compartments in the opposite direction. Inset: a “handshake” motif (discussed below), drives local reciprocal connectivity through postsynaptic terminals located on thin filopodia extending from two presynaptic boutons. **G** Electron mictoscopy cross-sections showing one synapse in each direction for the reciprocal connection between the LC10a neurons shown in F. **H** Illustration of a common motif that is analyzed in detail in Fig. 7: a postsynaptic terminal located on a thin filopodium extending from a multisynapse bouton (MSB).

Interestingly, visual projection neurons differ from both ALPNs and CX neurons in several additional respects. The fraction of connections occurring between neurons of the same cell type is more than threefold higher for visual projection neurons (Fig. 6A). At the same time, within-type visual projection neuron pairs show lower reciprocity and fewer synapses per connection than corresponding ALPN and CX pairs (Fig. 6C, D). Most ALPN types exhibit reciprocity rates exceeding 80%, whereas only a single LC type (LC10f; n = 4 neurons) reaches comparable levels. Thus, extensive same-type connectivity does not necessarily imply strong reciprocal connectivity.

Differences are also apparent in connectivity between neurons of different cell types (Supplementary Fig. 6–S3). ALPNs and visual projection neurons are both enriched in DD connections, whereas CX neurons are dominated by AD and AA connectivity. Together, these observations suggest that the organization of non-canonical connectivity cannot be explained solely by position along the sensory-to-motor axis and likely reflects additional circuit-specific computational requirements.

Examples of reciprocal connectivity in ALPNs and the visual projection neurons LC10a^63,64^ and LC31b^65^ are shown in Fig. 6E, F. Examination of synapse ultrastructure at AA and DD contact sites revealed a striking and recurrent morphological motif: postsynaptic terminals positioned directly on presynaptic boutons, often on thin filopodia extending from MSBs (Fig. 6E; analyzed in detail in Fig. 7). In some cases, neighboring boutons belonging to different neurons are reciprocally connected through such postsynaptic filopodia, forming a motif we term a “handshake” (Fig. 6F). Although the reciprocal synapses are located in close proximity, each synapse retains clear ultrastructural directionality, with distinct presynaptic specializations visible in both directions of the connection (Fig. 6G).

**Fig. 7:**
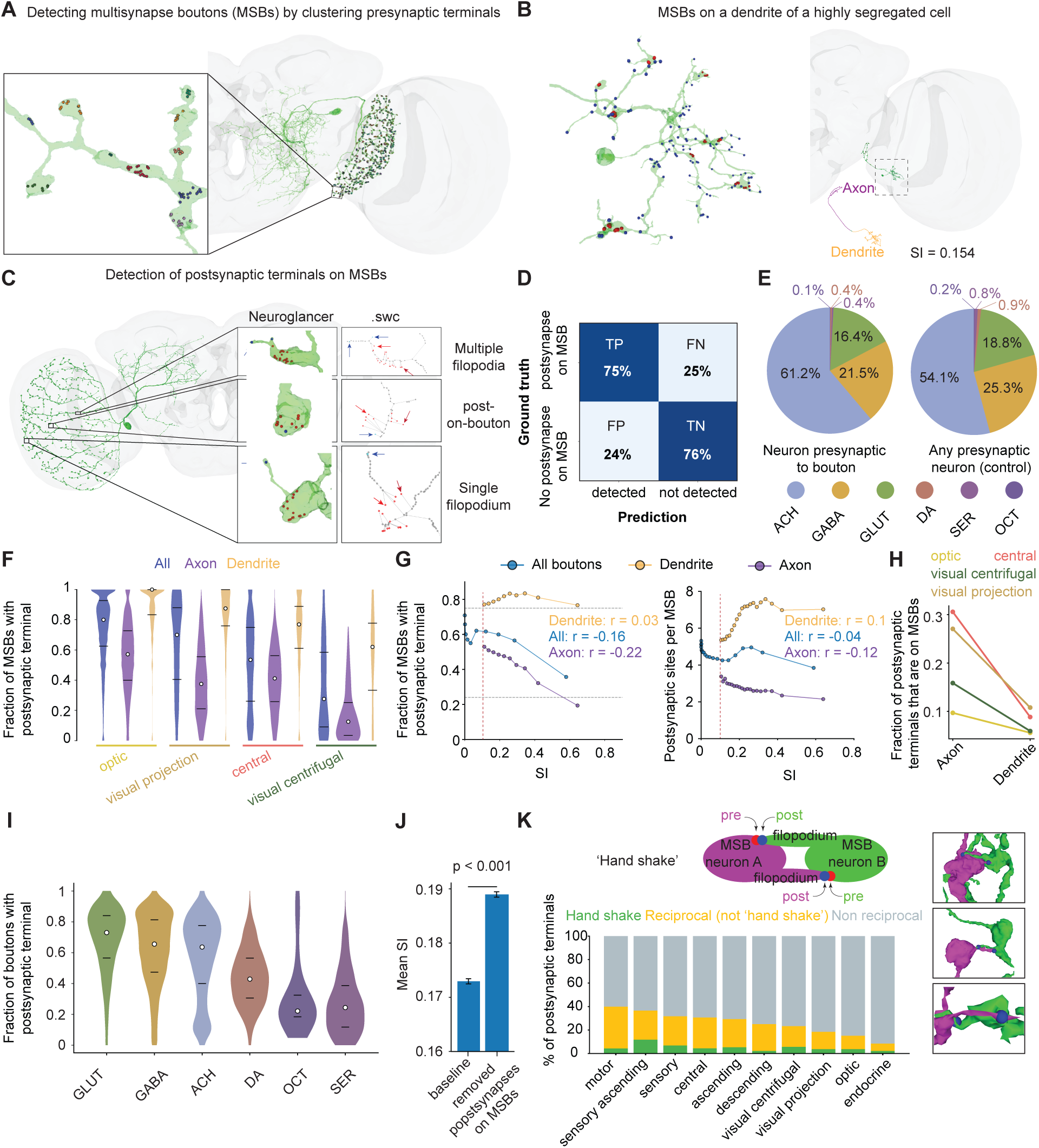
Postsynaptic terminals on MSBs contribute to mixed polarity and local recurrence. **A** An example neuron illustrating clustering of presynaptic terminals into putative multisynaptic boutons (MSBs). Presynaptic terminals were clustered using DBSCAN based on geodesic distances along the neuronal skeleton; each color denotes a distinct presynaptic cluster corresponding to an individual MSB. Neuron type: LT42 (visual centrifugal). **B** A polar neuron (segregation index [SI] = 0.15) exhibiting MSBs on its dendrite. The neuron is shown with its division into axonal and dendritic compartments, together with a close-up of the dendrite highlighting several distinct MSBs. **C** Filopodia on an example neuron (aMe17a1; see Table 2) illustrating three examples. Top: two postsynaptic terminals on a filopodium extending from an MSB and splitting into two branches. Middle: a single postsynaptic terminal located directly on an MSB. Bottom: a single postsynaptic terminal on a long thin filopodia extending from an MSB. Right: skeleton representations of the three examples, as extracted from the .swc file. Skeletons and pre-/post-synaptic terminals located along them were used to detect postsynaptic terminals on MSBs (see Methods). **D** Confusion matrix evaluating 200 pseudorandom cases (see Methods and Supplementary Fig. 7 - S3): 100 positive cases (one or more postsynaptic terminals on the MSB) and 100 negative cases (no postsynaptic terminals on the MSB). The model correctly identified 75/100 positive cases (true positives) and 76/100 negative cases (true negatives). **E** Distribution of presynaptic neurons across predicted neurotransmitter classes. Presynaptic neurons contacting postsynaptic terminals on MSBs were compared with the overall population of presynaptic neurons (control). Presynaptic neurons connected to postsynaptic terminals on MSBs were enriched for predicted cholinergic (ACH) neurons relative to the control population. **F** Fraction of MSBs containing one or more postsynaptic terminals for intrinsic neurons, separated by superclass and neuronal compartment, for neurons with a segregation index (SI) ≥ 0.1. The fraction was first calculated per neuron. White dots indicate medians and black lines indicate quartiles. **G** Left: fraction of MSBs containing one or more postsynaptic terminals as a function of SI. MSBs were divided into ten equal-sized bins according to the SI of the corresponding neuron, and the fraction of MSBs containing postsynaptic terminals was calculated for each bin. As in F, the analysis was restricted to intrinsic neurons with SI ≥ 0.1 and is shown separately for each superclass and compartment. Dashed horizontal lines represent the FP rate (0.24) and the (1 - FN) rate (0.75). These values correspond to the expected detector output if 0% or 100% of MSBs contained postsynaptic terminals, respectively. Right: mean number of postsynaptic terminals per MSB as a function of SI. MSBs were divided into 20 equal-sized bins according to the SI of the corresponding neuron, and the average number of postsynaptic terminals per MSB was calculated for each bin. For all correlations shown in G, p < 10⁻^30^. **H** Mean fraction of MSBs containing one or more postsynaptic terminals in axonal and dendritic compartments, calculated for neurons with SI ≥ 0.1 and shown separately for each intrinsic superclass as well as for all intrinsic neurons combined. **I** Fraction of MSBs containing one or more postsynaptic terminals for each predicted neurotransmitter class, using the same neurons analyzed in F–H. As in F, the fraction was first calculated per neuron. White circles indicate medians and black lines indicate quartiles. **J** Mean SI. Following targeted removal, SI increased significantly more than following random removal of an equal number of postsynaptic terminals (“baseline”), restricted to intrinsic superclasses but not limited to neurons with SI ≥ 0.1. Mean SI was first calculated per neuron. Following targeted removal, mean SI increased significantly relative to the baseline condition (Δ = 0.0160, 95% CI [0.0158, 0.0162]; paired t-test: p < 0.001, d = 0.50). **K** Top: schematic of a “handshake” connection between neuron A (purple) and neuron B (green), in which two MSBs, one belonging to each neuron, are reciprocally connected through postsynaptic terminals located on the two MSBs. Right: three example handshakes (only the postsynaptic terminals are shown). Bottom: postsynaptic terminals on MSBs were categorized as (1) participating in a handshake (green), (2) participating in a reciprocal connection between a pair of neurons, but not through a local handshake (yellow; see Supplementary Fig. 7 - S4), or (3) participating in neither, meaning that the neuron presynaptic to the postsynaptic terminal on the MSB is not reciprocally connected to the neuron containing that MSB. Neurons were assigned to superclasses according to the neuron containing the postsynaptic terminal on the MSB. For links to FlyWire examples, see Table 2.

Taken together, these examples illustrate that different circuit classes implement reciprocity through distinct combinations of canonical and non-canonical synaptic motifs. At the same time, they reveal a common ultrastructural feature—postsynaptic terminals located on MSBs—that may provide a general substrate for local recurrent connectivity. Therefore, we next asked how widespread postsynaptic terminals on MSBs are across the adult fly brain and whether they contribute systematically to mixed polarity and reciprocal connectivity.

### Postsynaptic terminals on boutons are a widespread substrate for non-canonical and reciprocal connectivity

Throughout the analyses above, we repeatedly observed postsynaptic terminals positioned on presynaptic boutons (Fig. 6E, H), motivating a systematic search for this motif across the FlyWire/FAFB connectome. We first identified MSBs by clustering presynaptic terminals using geodesic distances along neuronal skeletons (Fig. 7A; Supplementary Fig. 7–S1A), excluding cell types in which boutons could not be reliably separated (Supplementary Fig. 7–S1–S2 and Table 3; see Methods). Because our approach relies on clustering synapses rather than directly reconstructing bouton volumes, it is limited to MSBs, which are common throughout the *Drosophila* brain. To reduce false detections, we restricted the analysis to MSBs containing three or more presynaptic terminals. Notably, MSBs were detected not only on axons but also on dendrites, including dendrites of highly polarized neurons (Fig. 7B). We then identified postsynaptic terminals located in close proximity to these MSBs using neuronal skeletons represented as .swc files (Fig. 7C), enabling automated detection of postsynaptic terminals on MSBs throughout the brain (see Methods). Postsynaptic terminals associated with MSBs exhibit diverse morphologies: some are located directly on the bouton surface, whereas others are positioned on one or more filopodia extending from the bouton (Fig. 7C; Supplementary Fig. 7–S3A).

Because the detection algorithm relies on skeletal representations of neuronal arbors, it does not capture all postsynaptic terminals associated with boutons. Manual inspection of 200 pseudorandomly selected MSBs (100 positive and 100 negative cases; see Methods) revealed false-positive and false-negative rates of 24% and 25%, respectively (Fig. 7D). A typical false-positive case occurs when a postsynaptic terminal lies close to a bouton but is positioned along the main neurite, between neighboring boutons, rather than on the bouton surface or on an associated filopodium (Supplementary Fig. 7–S3B). Some such cases could reasonably be classified as bouton-associated by a human observer, suggesting that the estimated false-positive rate is conservative. Importantly, the fraction of MSBs carrying one or more detected postsynaptic terminals is highly reproducible both within cell types and between left–right homologs (Supplementary Fig. 7–S4A–B).

Postsynaptic terminals associated with presynaptic boutons have been reported previously in several mammalian brain regions, where they are often discussed in the context of presynaptic inhibition (see Discussion). To examine whether similar neurotransmitter-specific biases exist in the fly brain, we compared the predicted neurotransmitter identity of neurons presynaptic to postsynaptic terminals on MSBs with that of all presynaptic neurons in the dataset. Presynaptic partners of postsynaptic terminals on MSBs were enriched for cholinergic neurons relative to the overall population (Fig. 7E), indicating that this motif is not restricted to inhibitory circuits and may participate broadly in excitatory pathways as well. However, because acetylcholine can mediate either excitation or inhibition depending on receptor expression (see e.g.,^25^), the functional significance of this enrichment remains unclear.

Postsynaptic terminals on MSBs are remarkably widespread but vary substantially across neuronal classes. In some populations, the motif is extremely common. For example, approximately 80% of dendritic MSBs in optic neurons carry one or more postsynaptic terminals, whereas fewer than 30% of MSBs in visual centrifugal neurons do (Fig. 7F). Large differences are also observed across non-intrinsic superclasses (Supplementary Fig. 7–S4C), although these classes cannot be reliably subdivided into axonal and dendritic compartments because of incomplete reconstructions.

The prevalence of postsynaptic terminals on MSBs also strongly depends on local neuronal context. Across all intrinsic superclasses, dendritic MSBs are significantly more likely to carry postsynaptic terminals than axonal MSBs (Fig. 7F). Furthermore, the fraction of MSBs containing postsynaptic terminals depends on both neuronal compartment and overall neuronal polarity (Fig. 7G, left). On dendrites, this fraction approaches the upper bound imposed by the detector’s true-positive rate, suggesting that postsynaptic terminals may be present on nearly all dendritic MSBs. In contrast, on axons, the fraction decreases steadily as neuronal polarity increases, eventually approaching the detector’s false-positive baseline, indicating that postsynaptic terminals on axonal MSBs become rare in highly polarized neurons. These observations suggest that the probability of forming a postsynaptic terminal on an MSB depends on the local density of nearby postsynaptic sites, potentially reflecting shared developmental, structural, or signaling constraints.

Consistent with this interpretation, the average number of postsynaptic terminals associated with individual MSBs also varies systematically with neuronal polarity (Fig. 7G, right). As neurons become more polarized, dendritic MSBs tend to contain more postsynaptic terminals, whereas axonal MSBs contain fewer. Across all intrinsic superclasses, a substantially larger fraction of axonal postsynaptic terminals are associated with MSBs than dendritic postsynaptic terminals (Fig. 7H). For example, in central neurons, only approximately 10% of all dendritic postsynaptic terminals are associated with MSBs, compared with approximately 30% of axonal postsynaptic terminals. These values are likely underestimates because the analysis excludes boutons with only one or two presynaptic terminals.

The fraction of MSBs carrying postsynaptic terminals was relatively high and varied only modestly among the three major fast neurotransmitter classes: acetylcholine, GABA, and glutamate. By contrast, neurons predicted to release the major neuromodulators dopamine, octopamine, and serotonin showed substantially lower fractions. (Fig. 7I). Interestingly, however, neuromodulatory neurons are overrepresented among neurons presynaptic to postsynaptic terminals on MSBs (Fig. 7E), suggesting that this motif may provide a substrate for local modulation of synaptic output.

Although postsynaptic terminals on MSBs have only a modest effect on overall neuronal polarity (Fig. 7J), they are frequently associated with reciprocal connectivity. More than 25% of postsynaptic terminals on MSBs involve neuronal partners that are reciprocally connected to the neuron containing the bouton (Fig. 7K). A subset of these participates in highly localized bouton-to-bouton reciprocal interactions, which we term handshake motifs (Figs. 6F–H, 7K). Thus, postsynaptic terminals on MSBs are not merely anatomical curiosities but emerge as a fundamental anatomical motif linking mixed polarity, non-canonical connectivity, and local recurrent interactions.

Together, these analyses identify postsynaptic terminals on presynaptic boutons as a widespread and previously underappreciated circuit motif in the fly brain. This motif occurs across neuronal classes, is particularly enriched in regions and compartments exhibiting mixed polarity, and participates disproportionately in reciprocal connectivity. Although individual postsynaptic terminals on MSBs contribute only modestly to overall neuronal polarity, their collective prevalence indicates that they represent a substantial source of local input to presynaptic output structures.

More broadly, the widespread occurrence of postsynaptic terminals on boutons provides a concrete anatomical substrate for several of the organizational principles identified throughout this study. First, it offers a structural mechanism for mixed polarity by enabling input and output synapses to coexist within highly localized subcellular domains. Second, it supports non-canonical connectivity, including axo-axonic and dendrodendritic interactions. Third, it facilitates local reciprocal signaling, ranging from distributed reciprocal motifs to highly localized bouton-to-bouton handshake connections. The prevalence of this motif, together with the widespread occurrence of non-canonical and reciprocal connectivity across the connectome, suggests that mixed polarity in the adult *Drosophila* brain is not merely a consequence of incomplete segregation between axons and dendrites. Instead, it is supported by specific, recurrent anatomical motifs that appear across neuronal populations and brain regions, indicating that local recurrent interactions are a fundamental architectural feature of the fly brain.

## Discussion

Neurons with mixed presynaptic and postsynaptic domains have been described in numerous neural circuits across both vertebrates and invertebrates^23,26,29^. However, their prevalence and significance have remained difficult to assess at the whole-brain scale. Using the complete adult female *Drosophila* connectome, we show that mixed polarity is not restricted to a small number of specialized cell types but instead represents a widespread and highly structured feature of brain organization. Nearly half of intrinsic neurons exhibit low segregation indices; mixed polarity varies systematically across cell types, neurotransmitter classes, and brain regions; and these patterns are highly reproducible across hemispheres and among neurons of the same cell type. Together, these observations provide evidence against the idea that mixed polarity primarily reflects developmental immaturity or stochastic variation in neuronal organization. Instead, mixed polarity appears to be an integral component of circuit architecture.

Importantly, our results do not imply that mixed polarity is equally prevalent across all nervous systems. Although examples of mixed polarity and non-canonical connectivity have been reported in vertebrate circuits, including dendrodendritic interactions in the olfactory bulb and axo-axonic interactions in cortical and hippocampal networks^12,19,31,32,66^, the extent to which mixed polarity represents a major organizational feature of vertebrate connectomes remains unclear. Resolving this question will require systematic analyses of emerging large-scale datasets^67,68^.

A central result of this study is that synapse types are not distributed randomly across the brain. Rather, the prevalence of axo-dendritic (AD), axo-axonic (AA), dendrodendritic (DD), and dendro-axonic (DA) interactions can be predicted from simple properties of the participating neurons. Remarkably, a single axis of neuronal complexity derived from morphology and connectivity captures a substantial fraction of this organization. The first principal component explains 87% of the variance among eight morphological and connectivity features, indicating that these properties largely reflect a common underlying dimension ranging from small, weakly connected neurons to large, highly connected neurons. Synapse type can then be predicted from the positions of the pre- and post-synaptic neurons along this axis.

The resulting decision surface reveals two distinct organizational regimes. In one regime, dominated by optic neurons, interactions between neurons of similar complexity are primarily dendrodendritic. In the other, dominated by central and visual projection neurons, interactions between neurons of similar complexity are primarily axo-axonic. Independent observations support this distinction. Early sensory processing regions, including the optic lobe, antennal lobe, and AMMC, are enriched in DD connectivity, whereas more central regions, including the central complex and mushroom body output pathways, are enriched in AA connectivity^60–62^. Although these observations do not establish a strict processing hierarchy, they suggest that dendritic interactions may be preferentially associated with early sensory processing, whereas axonal interactions become increasingly prominent in downstream integrative circuits.

An additional and potentially more intriguing feature of the decision surface is the reversal of the directionality of canonical AD interactions across these regimes. Within the optic-neuron regime, canonical AD connections tend to link more complex neurons to simpler neurons, whereas in the central-brain regime, they tend to link simpler neurons to more complex neurons. The biological significance of this reversal remains unclear. One possibility is that neuronal complexity reflects different computational roles in different circuit architectures. Alternatively, the relationship may emerge from distinct developmental constraints acting on sensory and central circuits. Regardless of mechanism, the existence of a highly structured decision surface suggests that synapse type is itself an organizational property of neural circuits rather than merely a byproduct of neuronal geometry.

Our results further reveal a close relationship between mixed polarity and reciprocal connectivity. Reciprocal connections are a conserved feature of neural networks and are consistently overrepresented relative to random graph models across multiple nervous systems^45,48,49,60^. Previous work has described reciprocal motifs involving dendrites, axons, and mixed compartments in both vertebrate and invertebrate circuits^33,69–72^, but their relative prevalence has remained difficult to quantify at the whole-brain scale. We find that reciprocal connectivity is strongly enriched among neurons with mixed polarity and that non-canonical interactions dominate reciprocal motifs. Same-type reciprocal pairs are primarily connected through AA↔AA and DD↔DD motifs, whereas different-type pairs are dominated by complementary AD↔DA motifs. Purely canonical AD↔AD reciprocity is comparatively rare. These findings suggest that mixed polarity substantially expands the repertoire of recurrent architectures available to neural circuits.

The prominence of DD↔DD motifs in early sensory regions is particularly notable. In the fly, both the optic system and the antennal lobe contain extensive DD connectivity. The latter observation is particularly intriguing because the insect antennal lobe and the vertebrate olfactory bulb are among the clearest examples of convergent circuit organization across phyla, sharing glomerular architecture, extensive local interactions, and recurrent processing^73^. In vertebrates, DD reciprocity is a defining feature of olfactory bulb circuitry^19,29,74^. The enrichment of DD connectivity in the fly antennal lobe therefore raises the possibility that local dendritic interactions represent a conserved computational strategy for early olfactory processing, although testing this hypothesis will require direct comparative analyses.

At the subcellular level, we identify a widespread anatomical motif that may support many of these non-canonical interactions: postsynaptic terminals located directly on presynaptic boutons. Such arrangements have been described in individual circuits and are often discussed in the context of presynaptic inhibition or local synaptic modulation^75–77^. However, to our knowledge, their prevalence has not previously been quantified across an entire nervous system. We find that postsynaptic terminals associated with MSBs are widespread, vary systematically across neuronal classes and compartments, and frequently participate in reciprocal connectivity. Particularly striking are local handshake motifs, in which neighboring boutons belonging to different neurons are reciprocally connected through postsynaptic terminals located on the two boutons. These structures provide a direct anatomical substrate for highly localized recurrent interactions occurring at the scale of individual synaptic output sites.

The functional role of postsynaptic terminals on boutons remains unclear. One possibility is that they provide local feedback capable of modulating neurotransmitter release from nearby presynaptic sites. Alternatively, they may support compartment-specific computations that are difficult to achieve through conventional dendritic integration alone. The observed enrichment of cholinergic neurons among partners of bouton-associated postsynaptic terminals suggests that these interactions are not restricted to classical inhibitory mechanisms, although the functional consequences will depend on receptor composition and downstream signaling pathways. Indeed, acetylcholine can mediate either excitation or inhibition depending on receptor identity, as illustrated by muscarinic receptor-dependent inhibition in the mushroom body^25^. Determining whether bouton-associated postsynaptic terminals mediate excitation, inhibition, neuromodulation, or combinations therefore will require direct physiological investigation.

The emergence of new connectomic resources creates several opportunities to extend these findings. Within *Drosophila*, comparisons among larval^41^, adult female^42–44,78^, and recently completed male connectomes^79^ may reveal how mixed polarity and synapse-type organization change across development, sex, and individuals. Such analyses will help determine whether the architectural principles identified here are conserved or modified across brains that differ in size, behavior, and developmental history.

More broadly, the rapidly increasing size and number of vertebrate datasets, including large-scale mouse cortical reconstructions from MICrONS^67^, whole-brain zebrafish connectomes^68^, and emerging songbird connectomes^80^, may make it possible to determine whether comparable principles govern synapse-type organization in larger and evolutionarily distant nervous systems. Such comparisons could reveal whether the relationship between neuronal complexity, synapse type, and reciprocity reflects general constraints on neural circuit design or represents a specialization of compact insect brains.

Equally important will be understanding how these motifs arise. Mixed polarity and postsynaptic terminals on boutons are unlikely to be randomly distributed developmental outcomes. Instead, they probably reflect specific developmental programs that target pre- and post-synaptic specializations to particular subcellular domains. Future experiments combining connectomics, genetics, and synaptic labeling approaches (including emerging tools capable of simultaneously visualizing pre- and post-synaptic specializations at high resolution^81^) should make it possible to determine how these structures form, whether they are modified by experience, and how they contribute to computation and behavior.

In summary, our results suggest that mixed polarity is not simply a departure from the classical polarized neuron. Rather, it enables a diverse repertoire of synaptic interaction modes whose usage follows reproducible organizational principles across the brain. At the cellular level, these principles relate synapse type to neuronal complexity and circuit position. At the network level, they shape reciprocal connectivity and recurrent interactions. At the subcellular level, they are implemented through local motifs such as postsynaptic terminals on boutons. Together, these findings identify synapse type as a previously underappreciated organizational axis of neural circuit architecture and establish a framework for investigating how non-canonical interactions contribute to information processing across nervous systems.

## Methods

### Adult female brain

All analyses were performed using the FlyWire/FAFB consortium dataset (version v783; ^42–44^), comprising 139,255 proofread neurons and 80,215,790 synapses from the brain of a 7-day-old adult female *Drosophila melanogaster*^53^ (data snapshot available at Codex; https://codex.flywire.ai/). Unless otherwise stated, synapses were identified using the Princeton synapse detector^53^. In specific analyses, we additionally used the Buhmann synapse detector^54^. The detector, neuron set, and synapse set used for each figure panel are detailed in Table 1.

Neuron cell types, superclasses, and community annotations were obtained from the FlyWire annotation framework and downloaded from Codex in January 2026. Following the definitions of Schlegel et al.^42^, neurons were classified as intrinsic if all of their projections were contained within the brain. The intrinsic superclasses comprise central neurons, optic neurons, visual projection neurons, and visual centrifugal neurons. Analyses involving whole-cell properties, such as axon–dendrite compartmentalization using SFC or calculation of the SI, were restricted to intrinsic neurons or performed separately for intrinsic and non-intrinsic populations.

Neurotransmitter assignments were obtained from the FlyWire neurotransmitter classifier 68. Neurotransmitter-based analyses were restricted to the 107,761 intrinsic neurons with an assigned neurotransmitter label. Neuropil assignments were based on the FlyWire anatomical segmentation. Bilateral neuropils were merged into single regions, resulting in 43 neuropils for analysis (35 bilateral pairs and 8 unpaired midline neuropils).

Unless otherwise stated, synapses were excluded if either the presynaptic or postsynaptic site was not assigned to one of the 139,255 proofread neurons, or if the synapse connected a neuron to itself. The only exception was Supplementary Fig. 2–S1A, in which we explicitly tested the effect of including synapses involving unproofread segments (“twigs”).

### Adult skeletons

Neuron skeletons (SWC format) were obtained from the FlyWire Consortium v783 dataset and processed in Python using Navis/fafbseg (Navis is a Python library for the analysis and visualization of neuronal morphology; https://zenodo.org/records/8191725). Synapses were associated with neuronal skeletons using a modified workflow based on fafbseg.flywire.get_synapses with attach=True. A cleft-score threshold of ≥30 was applied. Synapses located more than 10,000 nm from the nearest skeleton node were excluded from further analysis.

### Larval SWC skeletons and synapses

For larval analyses, we used the published first-instar (*L1*) *Drosophila melanogaster* central nervous system connectome^41^, obtained from CATMAID. We analyzed the publicly available neuronal skeletons and synapse annotations without additional reconstruction or modification. Analyses were restricted to the 2,650 neurons for which both axonal and dendritic compartments were present in the dataset. SI (see below) was calculated using the same procedure as for the adult brain.

### Axonal and dendritic compartments

Axonal and dendritic compartments were identified using SFC^40^ as implemented in Navis (split_axon_dendrite, metric = synapse_flow_centrality). For each neuron, SFC was computed using the attached synapse set with reroot_soma = True and flow_thresh = 1. Under this stringent threshold (flow_thresh = 1), only nodes with the maximal SFC value were assigned to the intermediate (“linker”) compartment separating axonal and dendritic domains. This criterion minimizes fragmentation of the intermediate compartment and ensures a single transition region between axonal and dendritic compartments when a polarity split is present.

### Neural polarity and definition of mixed compartments

Neuronal polarity was quantified using the SI, which measures the degree of segregation between presynaptic and postsynaptic terminals across the axonal and dendritic compartments identified by SFC. SI ranges from 0 (no segregation; identical pre-/post-synaptic composition in axonal and dendritic compartments) to 1 (complete segregation). SI was computed for each neuron following the definition and procedure described by Schneider-Mizell et al.^40^ using the SFC-derived compartment assignments. A full description of the SI formulation is provided in the original study. SI values were highly consistent across synapse annotation methods, yielding similar results when calculated using either the Princeton synapse detector^53^ or the Buhmann synapse detector^54^ (Supplementary Fig. 2–S1A–C).

For analyses involving synapse type (AD, AA, DD, and DA), we restricted the dataset to neurons with SI ≥ 0.1. For these neurons, each synapse was classified according to the compartments containing its presynaptic and postsynaptic terminals, using A for axon and D for dendrite. This yielded four possible synapse types: AD, AA, DD, DA.

In addition to neuron-level polarity, some analyses required classification of individual compartments as mixed or non-mixed. To this end, axonal and dendritic compartments were analyzed separately. A compartment was classified as mixed when the fraction of canonical terminals in that compartment fell below a threshold determined by the corresponding SI value (Fig. 3C–E; Supplementary Fig. 3–S1A, S2A–B). At SI = 0.1, axonal compartments contained on average 71.0% presynaptic terminals and dendritic compartments contained on average 67.3% postsynaptic terminals (Supplementary Fig. 3–S1A). Therefore, axonal compartments with <71.0% presynaptic terminals and dendritic compartments with <67.3% postsynaptic terminals were classified as mixed.

### Predicting segregation index from morphology and connectivity features

SI was predicted from neuronal features using linear regression and random forest regression models implemented in scikit-learn with default hyperparameters (Fig. 2E). The eight predictor variables comprised four morphological features: size (volume), surface area, skeleton length, and number of leaves, and four connectivity features: number of input synapses, number of output synapses, number of input partners, and number of output partners.

Morphological features were obtained either from Codex (size, surface area, and skeleton length) or extracted directly from SWC files (number of leaves). Connectivity features were calculated from the Princeton synapse dataset and included the number of presynaptic terminals (output synapses), the number of postsynaptic terminals (input synapses), the number of output partners (distinct postsynaptic neurons), and the number of input partners (distinct presynaptic neurons). 10,000 randomly selected neurons from FlyWire were used for the prediction.

For each model, neurons were randomly split into training (80%) and test (20%) sets using a fixed random seed. Model performance was evaluated on the held-out test set using the coefficient of determination (R²). For random forest models, feature importance was quantified using the mean decrease in impurity (MDI), defined as the average reduction in prediction error attributable to splits on a given feature across all trees in the forest (Fig. 2E; Supplementary Fig. 2–S4B).

### Principal component analysis

PCA was applied to the eight neuronal features described above. Features were z-scored prior to decomposition. The weights of the first principal component (PC1) are shown in Fig. 2F. PC1 explained 87% of the total variance and was subsequently used as a low-dimensional measure of neuronal complexity (Figs. 2F-G, 4A–D).

### Predicting synapse type from morphology and connectivity features

Synapse type (AD, AA, DD, or DA) was predicted using four models (Fig. 4A, B), as follows. (i) Neuron identity model (“neurons”) – synapse type was predicted from the identities of the presynaptic and postsynaptic neurons. For each neuron pair, the predicted synapse type was the most common synapse type observed between that specific pair of neurons. Prediction accuracy is therefore limited by connection-type impurity (Fig. 3I). (ii) Cell-type model (“cell types”) – synapse type was predicted from the cell types of the presynaptic and postsynaptic neurons. For each presynaptic cell type–postsynaptic cell type combination, the predicted synapse type was the most common synapse type observed across all neuron pairs belonging to those cell types. (iii) Feature model (“features”) – synapse type was predicted using 16 neuronal features: 4 morphological and 4 connectivity features for the presynaptic neuron and the same 8 features for the postsynaptic neuron. (iv) PC1 model (“PC1”) – synapse type was predicted using only the first principal component (PC1) of the presynaptic and postsynaptic neurons (Figs. 2F, 4C–D). For machine-learning models (feature and PC1 models), classifiers were implemented in scikit-learn using default hyperparameters. To ensure balanced classification, the training dataset was subsampled to include all DA synapses and an equal number of synapses from each of the remaining synapse classes (AD, AA, and DD), yielding 541,126 synapses prior to train–test partitioning. Data were divided into training and test sets using an 80/20 stratified split to preserve class proportions in both sets. Model performance was evaluated on the held-out test set.

### Connection definition and connection purity

A directed connection from neuron A to neuron B was defined as the presence of at least one synapse with a presynaptic terminal on neuron A and a postsynaptic terminal on neuron B. No minimum synapse-count threshold was applied.

A connection was classified as pure if all of its synapses belonged to the same synapse type (Fig. 3I). For example, a connection containing both AA and DD synapses was considered non-pure, whereas a connection composed exclusively of AD synapses was considered pure.

For each synapse type, we quantified connection composition in two complementary ways. First, among all directed connections containing at least one synapse of a given type, we calculated the fraction of synapses belonging to each synapse type (Fig. 3I, left). Second, for the same set of connections, we calculated the fraction that also contained at least one synapse of each other synapse type (Supplementary Fig. 3–S2D).

Connection purity (Fig. 3I, right) was defined as the fraction of connections containing a given synapse type that were pure, i.e., composed exclusively of that synapse type.

### Reciprocity detection and reciprocal fraction

Two neurons were defined as reciprocally connected if at least one synapse was present in each direction (A→B and B→A). No minimum synapse-count threshold was imposed.

For each neuron, the reciprocal fraction was calculated as the number of reciprocal partners divided by the total number of unique synaptic partners (Fig. 5A).

Reciprocal neuron pairs were further characterized according to the dominant synapse type in each direction, defined as the synapse type contributing more than 50% of synapses in that direction (Fig. 5E–H; Supplementary Fig. 5–S1). Reciprocal pairs lacking a dominant synapse type in one or both directions were excluded from analyses requiring dominant-type classification (Fig. 5G, H). Reciprocal pairs were additionally classified according to whether the two neurons belonged to the same cell type and whether they were located in the same or opposite hemispheres.

### Synaptic strength bias toward reciprocal partners

For each neuron, we quantified whether reciprocal partners (bidirectionally connected neurons) received similar, stronger, or weaker synaptic investment than non-reciprocal partners (unidirectionally connected neurons). This was achieved by comparing the mean number of synapses per directed connection for reciprocal and non-reciprocal partners.

Specifically, for each neuron, we calculated (i) the mean number of synapses per non-reciprocal connection and (ii) the mean number of synapses per reciprocal connection. These quantities correspond to the x- and y-axes, respectively, in Fig. 5B. Points above the identity line indicate neurons that devote more synapses, on average, to reciprocal connections than to non-reciprocal connections, whereas points below the identity line indicate the opposite pattern. Neurons lacking either reciprocal or non-reciprocal partners were excluded from this analysis.

### Validation of DA synapses

To evaluate the possibility that some DA synapses identified by the Princeton detector reflect misassigned synapse direction, we manually examined 200 randomly selected synapses (50 from each synapse class: AD, AA, DD, and DA). The observer was blinded both to the synapse classification and to the detector-assigned synapse direction. Synapse direction was determined using ultrastructural markers of presynaptic and postsynaptic specializations, specifically T-bars and synaptic vesicles for the presynaptic terminal and postsynaptic densities for the postsynaptic terminal.

For 70% of the DA synapses, the manually assigned direction agreed with the direction assigned by the Princeton detector. Agreement was higher for the other synapse classes: 100% (AD), 92% (AA), and 98% (DD). Thus, DA synapses showed substantially lower direction-assignment agreement than the other synapse classes.

Assuming that direction-assignment errors occur independently across synapses, the observed disagreement rate for DA synapses (30%) can be used to estimate the probability that a putative DA connection contains no true DA synapses. Under this assumption, 30% of neuron pairs containing a single detected DA synapse would be expected to contain no true DA synapse. For pairs containing two or three detected DA synapses, the corresponding probabilities decrease to 9.0% and 2.7%, respectively.

Given the observed distribution of DA synapse counts across neuron pairs, and assuming independent direction-assignment errors, we estimate that 79.3% of pairs containing one or more detected DA synapses contain at least one true DA synapse.

### Detecting putative MSBs by clustering of presynaptic terminals

For each neuron, MSBs were identified by clustering presynapses based on geodesic distances along the neuronal skeleton using DBSCAN. Neurons were represented as .swc files, in which each neuron forms a tree of interconnected nodes that describe its morphology. Each synapse is associated with a single skeleton node. To compute distances between synapses, we extracted geodesic distances, defined as the shortest path along the neuronal tree (rather than Euclidean distance), using navis.geodesic_matrix.

Pairwise geodesic distances between nodes having associated presynaptic terminals were computed and further processed for clustering using DBSCAN (eps = 900, min_samples = 3). Due to occasional inconsistencies in skeleton topology relative to the true neuronal morphology, some presynapses were either incorrectly left unclustered or split into multiple clusters within a single bouton. To correct this, any non-clustered presynapse located within 450 nm (Euclidean distance) of a cluster was assigned to that cluster, and then clusters separated by less than 450 nm (minimum Euclidean distance between cluster centers) were merged.

Certain cell types and subclasses were known a priori to lack typical bouton organization, often forming large continuous aggregates of presynapses or other structures that do not correspond to discrete boutons. These cell types were therefore excluded before clustering. Excluded groups included the visual receptor neurons R1–R8; the optic neurons of types L1–L5, Tm1–Tm4, Tm5a, Tm5f, Tm6, Tm9, Tm21, Mi1, Mi4, T1, and T3; and all neurons belonging to the “uniglomerular” subclass.

In addition to these predefined exclusions, further atypical cell types were identified through a post hoc quality-control analysis of clustering results. For each neuron, we calculated the root-mean-square (RMS) distance of presynapses from their assigned cluster centroid:

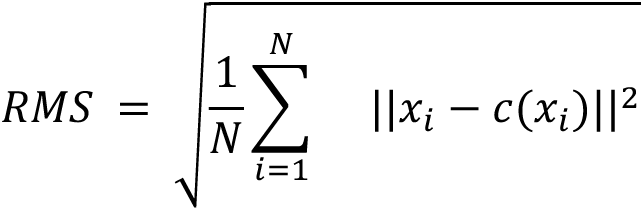

where *x_i_* is the position of presynapse *i*, *c*(*x_i_*) is the centroid of the cluster to which that presynapse was assigned, and *N* is the total number of clustered presynapses for the neuron.

Neurons with RMS values greater than 850 nm were manually inspected. These neurons frequently exhibited unusually large and spatially dispersed clusters that were inconsistent with bouton-like organization. Examination of their corresponding cell types revealed several additional neuronal classes that had not been identified during the initial exclusion stage. Notably, some of these classes were closely related to cell types that had already been excluded a priori, suggesting similar underlying morphological organization.

As an additional diagnostic metric, we calculated the mean pairwise Euclidean distance between cluster centroids within each neuron. For a neuron containing *K* clusters with centroid coordinates *c*_1_, . . . , *c_k_* this metric was defined as:

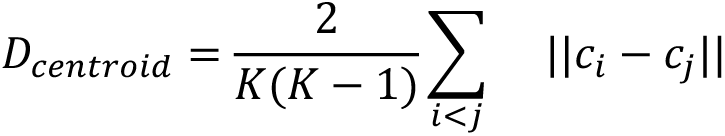

where ||*c_i_* – *c_j_*|| denotes the Euclidean distance between centroids *i* and *j*. This quantity corresponds to the average of all pairwise distances between cluster centroids within a neuron. Unlike the RMS metric, mean centroid distance did not identify neurons with atypical bouton organization. Instead, it highlighted a small set of neurons with exceptionally large inter-cluster separations. Manual inspection revealed that most of these neurons belonged to the Bilateral class. These neurons were not excluded from the analysis, as their large centroid distances reflected genuine anatomical organization, with boutons distributed across both the left and right optic lobes rather than abnormal clustering behavior. Thus, mean centroid distance served primarily as a diagnostic measure for identifying neurons with unusually broad spatial distributions of boutons, whereas the RMS metric was more effective for detecting neurons with large and spatially dispersed clusters that deviated from typical bouton organization.

After applying all exclusion criteria - including predefined cell-type exclusions, the uniglomerular subclass exclusion, and the RMS-based post hoc exclusions - a total of 53,561 neurons were removed from the original dataset of 139,255 (for the full list, see Table 3).

### Detecting postsynaptic terminals on MSBs

Postsynaptic terminals on MSBs were detected using a three-step procedure based on neuronal skeletons (.swc files) and synapse annotations.

Step 1: Detection of candidate postsynaptic terminals.

For each presynaptic cluster (putative MSB), a centroid node was identified based on the coordinates of the presynaptic terminals belonging to the cluster. Candidate postsynaptic terminals were then identified as all postsynaptic terminals located within a geodesic distance of 5,500 nm from the centroid node. This threshold was chosen empirically to favor sensitivity over specificity, as false-positive candidates are removed in subsequent filtering steps.

Step 2: Exclusion of terminals associated with neighboring presynaptic sites.

For each candidate postsynaptic terminal identified in Step 1, the geodesic distance to the nearest non-clustered presynaptic terminal was compared with the distance to the nearest MSB centroid. Candidates whose nearest non-clustered presynaptic terminal was closer than the nearest MSB centroid were excluded.

Step 3: Skeleton-based filtering.

For each remaining candidate, the path from the postsynaptic terminal toward the corresponding MSB centroid was defined as the inward direction. All other branches emerging from the candidate node were defined as outward directions. The neuronal skeleton was then traversed from the postsynaptic terminal along each outward branch. If a presynaptic terminal was encountered along any outward path, the candidate was excluded. The remaining candidates were classified as postsynaptic terminals on MSBs.

Steps 2 and 3 were designed to remove postsynaptic terminals located along the main neuronal process and retain only terminals that were spatially associated with a single MSB. Visual inspection indicated that most detected postsynaptic terminals were located on short filopodia extending from boutons. However, because the analysis relied on skeleton geometry rather than full volumetric reconstructions, we could not distinguish between postsynaptic terminals located directly on the bouton surface and those located on filopodia extending from the bouton.

### Validation of postsynapse-on-MSB detector

Detector performance was evaluated by manual inspection of 200 MSBs. To provide a stringent test of detector performance, candidate MSBs were sampled randomly under the constraint that at least one postsynaptic terminal was located within 6 µm of the MSB centroid. This criterion ensured that all evaluated MSBs contained a nearby postsynaptic terminal and therefore represented challenging classification cases.

A manual observer, blinded to the detector’s predictions, examined the sampled MSBs and classified each as positive (containing one or more postsynaptic terminals on the MSB) or negative. Annotation continued until 100 manually identified positive MSBs and 100 manually identified negative MSBs had been obtained. These manual annotations were treated as ground truth and compared with the detector’s predictions.

Based on this analysis, the detector exhibited false-positive and false-negative rates of 24% and 25%, respectively (Fig. 7D; Supp. Fig. 7–S3; Table 4).

### Statistical analyses

All statistical analyses were performed in Python 3 using SciPy, NumPy, scikit-posthocs, and scikit-learn. Normality was assessed using the Shapiro–Wilk test together with a median-based omnibus normality test, and homogeneity of variances was assessed using Levene’s test. Because these assumptions were frequently violated, group comparisons were generally performed using the Kruskal–Wallis test. When omnibus tests were significant, post-hoc pairwise comparisons were performed using Dunn’s test with Bonferroni correction. Effect sizes were quantified using η² for Kruskal–Wallis tests and rank-biserial correlation for pairwise comparisons (e.g., Fig. 2B, C).

For analyses in which residuals were approximately normally distributed but variances were unequal, Welch’s ANOVA was used. Post-hoc comparisons were performed using Welch’s t-tests with Bonferroni correction, and effect sizes were quantified using Hedges’ g (e.g., Fig. 1D).

Associations between continuous variables were quantified using Pearson correlation coefficients. Correlations were interpreted primarily based on effect size rather than statistical significance, given the large sample sizes used throughout the study. All statistical tests were two-tailed unless stated otherwise.

### Visualization and figure generation

Figures were generated in Python using Matplotlib and Seaborn. This included the violin plots, kernel density estimates, heatmaps, scatter plots, cumulative distribution plots, and confusion matrices.

Neuron morphologies and synaptic locations were visualized using Navis for three-dimensional skeleton rendering, FlyWire Neuroglancer for mesh-based visualization, and Codex point-cloud coordinates for synapse overlays.

Electron microscopy images and Neuroglancer visualizations (e.g., Fig. 1A) were obtained as screenshots from the corresponding platforms. Links to all FlyWire examples are provided in Table 2.

### Software environment

All analyses were performed in Python 3.12.4 using the packages Navis, CaveClient, NumPy, Pandas, SciPy, Matplotlib, Seaborn, scikit-learn, and scikit-posthocs. Computations were executed locally on Windows workstations equipped with 32 GB RAM.

The code used for data processing, analysis, and figure generation is available at: https://github.com/deutschlab/mixed-polarity-paper

## Acknowledgments

We are grateful to Greg Jefferis, Philipp Schlegel, Marion Silies, and Uri Hershberg for insightful discussions, thoughtful feedback, and constructive comments on the manuscript, which substantially improved the presentation of this work. This work was supported by the Zuckerman STEM Leadership Program.

## Tables

Table 1 Statistics

Table 2 Neuroglancer/FlyWire links

Table 3 Cell types excluded from MSB clustering

Table 4 Validation of postsynapse-on-MSB detection

## References

1. Cajal, S. R. y. Cajal’s Histology of the Nervous System of Man and Vertebrates. (Oxford University Press, New York, NY, 1995).

2. Ramón y Cajal, S. Textura del sistema nervioso del hombre y de los vertebrados : estudios sobre el plan estructural y composición histológica de los centros nerviosos adicionados de consideraciones fisiológicas fundadas en los nuevos descubrimientos. Volumen I. *Madrid : Nicolás Moya* https://digibug.ugr.es/handle/10481/69713 (1899).

3. Kandel, E., Koester, J. D., Mack, S. H. & Siegelbaum, S. Principles of Neural Science, Sixth Edition. (McGraw-Hill Education, Columbus, OH, 2021).

4. Shepherd, G. M. Foundations of the Neuron Doctrine. (Oxford University Press, New York, NY, 2016).

5. Rolls, M. M. & Jegla, T. J. Neuronal polarity: an evolutionary perspective. J Exp Biol 218, 572–580 (2015).

6. Takano, T., Xu, C., Funahashi, Y., Namba, T. & Kaibuchi, K. Neuronal polarization. Development 142, 2088–2093 (2015).

7. Goaillard, J.-M., Moubarak, E., Tapia, M. & Tell, F. Diversity of axonal and dendritic contributions to neuronal output. Front. Cell. Neurosci. 13, 570 (2019).

8. Barnes, A. P. & Polleux, F. Establishment of axon-dendrite polarity in developing neurons. Annu. Rev. Neurosci. 32, 347–381 (2009).

9. Kidd, M. Electron microscopy of the inner plexiform layer of the retina in the cat and the pigeon. J Anat 96, 179–187 (1962).

10. Colonnier, M. & Guillery, R. W. Synaptic organization in the lateral geniculate nucleus of the monkey. Z. Zellforsch. Mikrosk. Anat. 62, 333–355 (1964).

11. Hirata, Y. SOME OBSERVATIONS ON THE FINE STRUCTURE OF THE SYNAPSES IN THE OLFACTORY BULB OF THE MOUSE, WITH PARTICULAR REFERENCE TO THE ATYPICAL SYNAPTIC CONFIGURATIONS. Arch Histol Jpn 24, 293–302 (1964).

12. Rall, W. & Shepherd, G. M. Theoretical reconstruction of field potentials and dendrodendritic synaptic interactions in olfactory bulb (1968). in The Theoretical Foundation of Dendritic Function 173–204 (The MIT Press, 1994).

13. Yuste, R. & Tank, D. W. Dendritic integration in mammalian neurons, a century after Cajal. Neuron 16, 701–716 (1996).

14. Grimes, W. N., Zhang, J., Graydon, C. W., Kachar, B. & Diamond, J. S. Retinal parallel processors: more than 100 independent microcircuits operate within a single interneuron. Neuron 65, 873–885 (2010).

15. Hausselt, S. E., Euler, T., Detwiler, P. B. & Denk, W. A dendrite-autonomous mechanism for direction selectivity in retinal starburst amacrine cells. PLoS Biol. 5, e185 (2007).

16. Carden, W. B. & Bickford, M. E. Synaptic inputs of class III and class V interneurons in the cat pulvinar nucleus: differential integration of RS and RL inputs. Vis. Neurosci. 19, 51–59 (2002).

17. Famiglietti, E. V., Jr. Dendro-dendritic synapses in the lateral geniculate nucleus of the cat. Brain Res. 20, 181–191 (1970).

18. Morest, D. K. Synaptic relationships of Golgi type II cells in the medial geniculate body of the cat. J. Comp. Neurol. 162, 157–193 (1975).

19. Rall, W., Shepherd, G. M., Reese, T. S. & Brightman, M. W. Dendrodendritic synaptic pathway for inhibition in the olfactory bulb (1966). in The Theoretical Foundation of Dendritic Function 160–172 (The MIT Press, 1994).

20. Christiansen, F. et al. Presynapses in Kenyon cell dendrites in the mushroom body calyx of Drosophila. J. Neurosci. 31, 9696–9707 (2011).

21. Gray, E. G. Electron microscopy of excitatory and inhibitory synapses: a brief review. Prog. Brain Res. 31, 141–155 (1969).

22. White, J. G., Southgate, E., Thomson, J. N. & Brenner, S. The structure of the nervous system of the nematode Caenorhabditis elegans. Philos. Trans. R. Soc. Lond. B Biol. Sci. 314, 1–340 (1986).

23. Morgan, J. L. & Lichtman, J. W. An individual interneuron participates in many kinds of inhibition and innervates much of the mouse visual thalamus. Neuron 106, 468–481.e2 (2020).

24. Haag, J. & Borst, A. Dendro-dendritic interactions between motion-sensitive large-field neurons in the fly. J Neurosci 22, 3227–3233 (2002).

25. Manoim, J. E., Davidson, A. M., Weiss, S., Hige, T. & Parnas, M. Lateral axonal modulation is required for stimulus-specific olfactory conditioning in Drosophila. Curr. Biol. 32, 4438–4450.e5 (2022).

26. Schneider-Mizell, C. M. et al. Structure and function of axo-axonic inhibition. Elife 10, (2021).

27. Bates, A. S. et al. Complete connectomic reconstruction of olfactory projection neurons in the fly brain. Curr. Biol. 30, 3183–3199.e6 (2020).

28. DeFelipe, J. Brain plasticity and mental processes: Cajal again. Nat Rev Neurosci 7, 811–817 (2006).

29. Shepherd, G. M. Symposium overview and historical perspective: dendrodendritic synapses: past, present, and future. Ann. N. Y. Acad. Sci. 1170, 215–223 (2009).

30. Ramón y Cajal, S. Histologie du système nerveux de l’homme & des vertébrés: Cervelet, cerveau moyen, rétine, couche optique, corps strié, écorce cérébrale générale & régionale, grand sympathique. (1911).

31. Bartel, D. L., Rela, L., Hsieh, L. & Greer, C. A. Dendrodendritic synapses in the mouse olfactory bulb external plexiform layer. J. Comp. Neurol. 523, Spc1–Spc1 (2015).

32. Dudok, B. et al. Recruitment and inhibitory action of hippocampal axo-axonic cells during behavior. Neuron 109, 3838–3850.e8 (2021).

33. Boeckh, J. & Tolbert, L. P. Synaptic organization and development of the antennal lobe in insects. Microscopy Research and Technique 24, 260–280 (1993).

34. Sun, X. J., Tolbert, L. P. & Hildebrand, J. G. Synaptic organization of the uniglomerular projection neurons of the antennal lobe of the moth Manduca sexta: A laser scanning confocal and electron microscopic study. Journal of Comparative Neurology 379, 2–20 (1997).

35. Manoim-Wolkovitz, J. E. et al. Nonlinear high-activity neuronal excitation enhances odor discrimination. Curr. Biol. 35, 1521–1538.e5 (2025).

36. McGeer, P. L., McGeer, E. G. & Innanen, V. T. Dendro axonic transmission. I. Evidence from receptor binding of dopaminergic and cholinergic agents. Brain Res. 169, 433–441 (1979).

37. Hattori, T., McGeer, P. L. & McGeer, E. G. Dendro axonic neurotransmission. II. Morphological sites for the synthesis, binding and release of neurotransmitters in dopaminergic dendrites in the substantia nigra and cholinergic dendrites in the neostriatum. Brain Res. 170, 71–83 (1979).

38. Zhang, Y. et al. Identifying local and descending inputs for primary sensory neurons. J. Clin. Invest. 125, 3782–3794 (2015).

39. Duncan, D. & Morales, R. Relative numbers of several types of synaptic connections in the substantia gelatinosa of the cat spinal cord. J. Comp. Neurol. 182, 601–610 (1978).

40. Schneider-Mizell, C. M. et al. Quantitative neuroanatomy for connectomics in Drosophila. Elife 5, (2016).

41. Winding, M. et al. The connectome of an insect brain. Science 379, eadd9330 (2023).

42. Dorkenwald, S. et al. Neuronal wiring diagram of an adult brain. Nature 634, 124–138 (2024).

43. Schlegel, P. et al. Whole-brain annotation and multi-connectome cell typing of Drosophila. Nature 634, 139–152 (2024).

44. Zheng, Z. et al. A Complete Electron Microscopy Volume of the Brain of Adult Drosophila melanogaster. Cell 174, 730–743.e22 (2018).

45. Lin, A. et al. Network statistics of the whole-brain connectome of Drosophila. Nature 634, 153–165 (2024).

46. Gilbert, E. N. Random Graphs. Ann. Math. Stat. 30, 1141–1144 (1959).

47. Milo, R., Kashtan, N., Itzkovitz, S., Newman, M. E. J. & Alon, U. On the uniform generation of random graphs with prescribed degree sequences. arXiv [cond-mat.stat-mech] (2003).

48. Varshney, L. R., Chen, B. L., Paniagua, E., Hall, D. H. & Chklovskii, D. B. Structural properties of the Caenorhabditis elegans neuronal network. PLoS Comput. Biol. 7, e1001066 (2011).

49. Cook, S. J. et al. Whole-animal connectomes of both Caenorhabditis elegans sexes. Nature 571, 63–71 (2019).

50. Yang, R. et al. Cyclic structure with cellular precision in a vertebrate sensorimotor neural circuit. Curr. Biol. 33, 2340–2349.e3 (2023).

51. Turner, N. L. et al. Reconstruction of neocortex: Organelles, compartments, cells, circuits, and activity. Cell 185, 1082–1100.e24 (2022).

52. Perin, R., Berger, T. K. & Markram, H. A synaptic organizing principle for cortical neuronal groups. Proc. Natl. Acad. Sci. U. S. A. 108, 5419–5424 (2011).

53. 53. Yu, S.-C., et al. New synapse detection in the whole-brain connectome of *Drosophila*. bioRxiv (2025) doi:10.1101/2025.07.11.664377.

54. Buhmann, J. et al. Automatic detection of synaptic partners in a whole-brain Drosophila electron microscopy data set. Nat. Methods 18, 771–774 (2021).

55. Newman, M. Networks. (Oxford University Press, London, England, 2010).

56. Braun, J., Hurtak, F., Wang-Chen, S. & Ramdya, P. Descending networks transform command signals into population motor control. Nature 630, 686–694 (2024).

57. Meier, M. & Borst, A. Extreme compartmentalization in a Drosophila amacrine cell. Curr. Biol. 29, 1545–1550.e2 (2019).

58. Nern, A. et al. Connectome-driven neural inventory of a complete visual system. Nature (2025) doi:10.1038/s41586-025-08746-0.

59. Seung, H. S. Multi-output computation by single neuron biophysics in a visual system. Neuroscience (2025).

60. Hulse, B. K. et al. A connectome of the Drosophila central complex reveals network motifs suitable for flexible navigation and context-dependent action selection. Elife 10, e66039 (2021).

61. Gruber, L. et al. The unique synaptic circuitry of specialized olfactory glomeruli in Drosophila melanogaster. Elife 12, (2025).

62. Li, F. et al. The connectome of the adult Drosophila mushroom body provides insights into function. Elife 9, e62576 (2020).

63. Ribeiro, I. M. A. et al. Visual Projection Neurons Mediating Directed Courtship in Drosophila. Cell 174, 607–621.e18 (2018).

64. Hindmarsh Sten, T., Li, R., Otopalik, A. & Ruta, V. Sexual arousal gates visual processing during Drosophila courtship. Nature 595, 549–553 (2021).

65. 65. Deutsch, D., et al. Sexually-dimorphic neurons in the Drosophila whole-brain connectome. *bioRxivorg* (2025) doi:10.1101/2025.06.10.658788.

66. Cover, K. K. & Mathur, B. N. Axo-axonic synapses: Diversity in neural circuit function. J. Comp. Neurol. 529, 2391–2401 (2021).

67. MICrONS Consortium. Functional connectomics spanning multiple areas of mouse visual cortex. Nature 640, 435–447 (2025).

68. Svara, F. et al. Automated synapse-level reconstruction of neural circuits in the larval zebrafish brain. Nat. Methods 19, 1357–1366 (2022).

69. Sloper, J. J. & Powell, T. P. Dendro-dendritic and reciprocal synapses in the primate motor cortex. Proc. R. Soc. Lond. B Biol. Sci. 203, 23–38 (1978).

70. 70. Gilbert, E. T., et al. Reciprocal interactions between CA1 pyramidal and axo-axonic cells control sharp wave-ripple events. bioRxivorg (2025) doi:10.1101/2024.07.02.601726.

71. Cervantes-Sandoval, I., Phan, A., Chakraborty, M. & Davis, R. L. Reciprocal synapses between mushroom body and dopamine neurons form a positive feedback loop required for learning. Elife 6, e23789 (2017).

72. Isaacson, J. S. & Strowbridge, B. W. Olfactory reciprocal synapses: dendritic signaling in the CNS. Neuron 20, 749–761 (1998).

73. Laurent, G. Olfactory network dynamics and the coding of multidimensional signals. Nat. Rev. Neurosci. 3, 884–895 (2002).

74. Bartel, D. L., Rela, L., Hsieh, L. & Greer, C. A. Dendrodendritic synapses in the mouse olfactory bulb external plexiform layer: Dendrodendritic Synapses of Mouse EPL. J. Comp. Neurol. 523, 1145–1161 (2015).

75. Rudomin, P. & Schmidt, R. F. Presynaptic inhibition in the vertebrate spinal cord revisited. Exp. Brain Res. 129, 1–37 (1999).

76. Thompson, S. M., Capogna, M. & Scanziani, M. Presynaptic inhibition and facilitation of transmitter release in the hippocampus. in Presynaptic Inhibition and Neural Control 95–110 (Oxford University PressNew York, NY, 1997).

77. Guo, D. & Hu, J. Spinal presynaptic inhibition in pain control. Neuroscience 283, 95–106 (2014).

78. 78. Bates, A. S., et al. Distributed control circuits across a brain-and-cord connectome. bioRxivorg (2025) doi:10.1101/2025.07.31.667571.

79. Berg, S. et al. Sexual dimorphism in the complete connectome of the Drosophila male central nervous system. bioRxivorg (2025) doi:10.1101/2025.10.09.680999.

80. Rother, A., Januszewski, M., Jain, V., Fee, M. S. & Kornfeld, J. The songbird basal ganglia connectome. bioRxivorg (2025) doi:10.1101/2025.10.25.684569.

81. Parisi, M. J., Aimino, M. A. & Mosca, T. J. A conditional strategy for cell-type specific labeling of endogenous excitatory synapses in Drosophila reveals subsynaptic architecture. bioRxiv 2022.10.17.510548 (2022) doi:10.1101/2022.10.17.510548.

