## Supplemental Figures for "Organizational principles governing synapse types in a whole-brain connectome"

### Supplementary figures

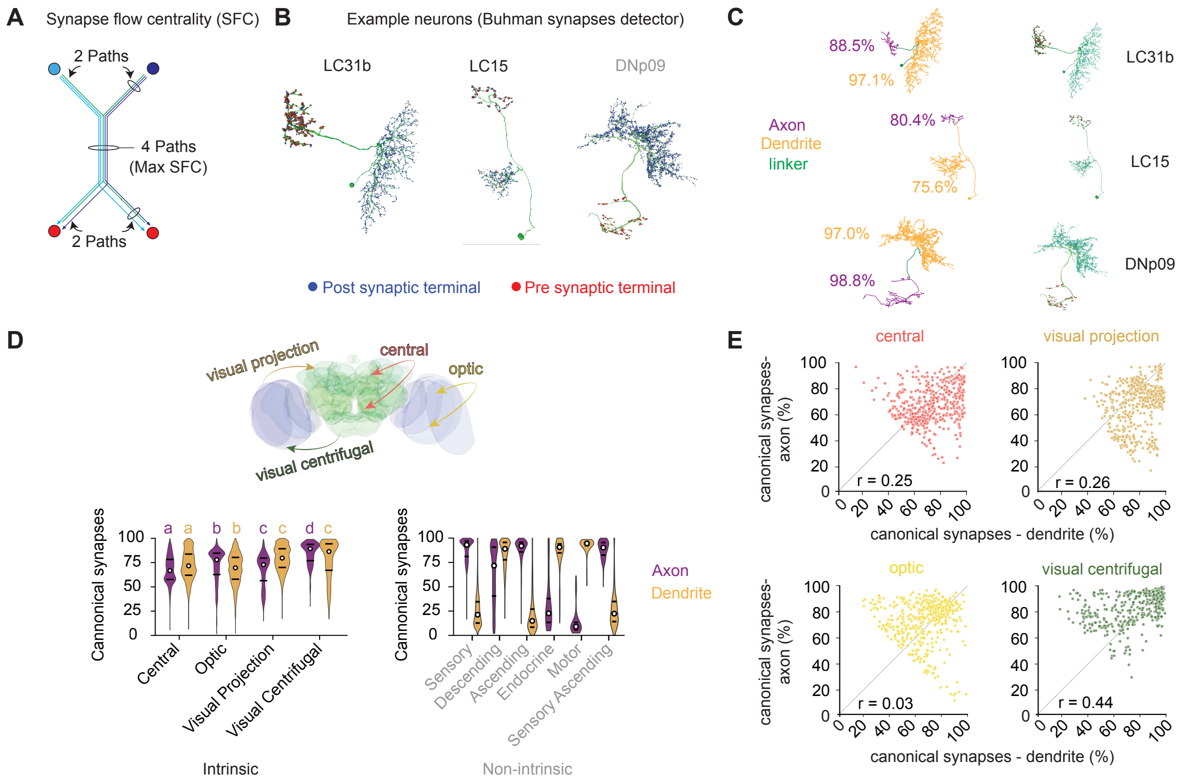


**Supp Fig. 1 - S1. Synaptic compartmentalization and canonicality. A** Schematic of 'Synapse Flow Centrality' (SFC) calculation. 'Max SFC' is defined as the largest number of paths that connect presynaptic and postsynaptic terminals across the skeleton. Max SFC is used to partition the neuron into axon and dendrite. **B** Neuroglancer renderings of three visibly distinguishable polarized neurons (LC31b, LC15, DNp09; same as Fig. 1B) showing skeletons (green), presynaptic terminals (red), and postsynaptic terminals (blue) using the 'Buhmann' synapse detector[1]. **C** Example neurons after the SFC process, with axonal (purple) and dendritic (orange) regions and their canonical synapse percentages per compartment, calculated as the ratio of canonical to total synapses using the Buhmann detection data set. Results are similar to those obtained using the 'Princeton synapses' (Fig. 1C). **D** Violin plots of axonal (purple) and dendritic (orange) canonicality per superclass. Left: intrinsic superclasses - central (32,226 cells), optic (76,818 cells), visual projection (8,008 cells), and visual centrifugal (518 cells). Right: non-intrinsic superclasses - sensory (16,106), descending (1,296), ascending (1,734), endocrine (78), motor (110), and sensory ascending (591). White dots denote medians and black bars represent interquartile ranges. For intrinsic neurons, Because variances were unequal across groups, we used Welch's ANOVA followed by Games–Howell post-hoc comparisons. Axonal canonicality differed across super-classes, $F(3, 2468.50) = 864.02$, $p < 0.001$, $\eta^2p = 0.016$; all pairwise comparisons were significant except central versus visual projection neurons ($p = 0.999$). Dendritic canonicality also differed across super-classes, $F(3, 2447.49) = 1756.96$, $p < 0.001$, $\eta^2p = 0.035$; all pairwise comparisons were significant except visual centrifugal versus visual projection neurons (p = 0.961). A significant compartment × super-class interaction, $F(3, 117566) = 1794.47$, $p < 0.001$, $\eta^2p = 0.044$, reflected opposite compartmental patterns across super-classes: optic and visual centrifugal neurons showed higher axonal than dendritic canonicality, whereas central and visual projection neurons showed higher dendritic than axonal canonicality. Inset: schematic classification of intrinsic neuron superclasses by neuropil localization. Green indicates central neuropils; blue indicates optic neuropils. Central neurons form inputs and outputs within central neuropils. Optic neurons arborize and form inputs and outputs within optic neuropils. Visual projection neurons receive inputs in optic neuropils and send outputs to central neuropils. Visual centrifugal neurons receive inputs in central neuropils and send outputs to optic neuropils. **E** Pearson correlation, comparing dendritic and axonal canonicality within each superclass.

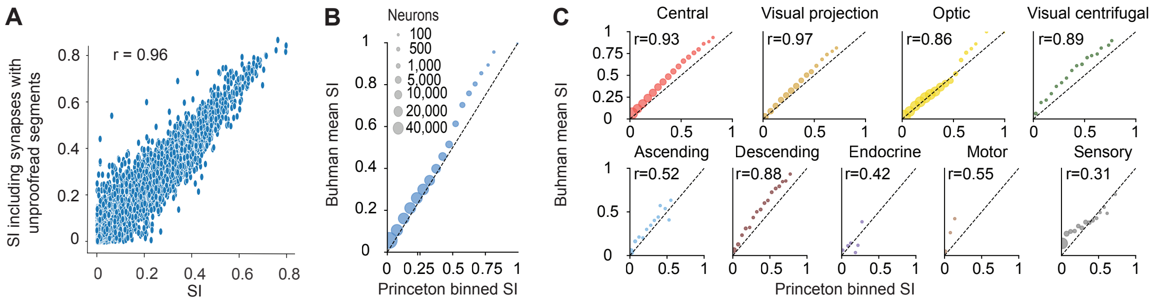


**Supp Fig. 2 - S1** **A** Comparison of SI values calculated using only synapses between proofread neurons in FlyWire/FAFB v783 (as in the remainder of the paper; see Methods) versus SI values calculated using all presynaptic and postsynaptic terminals of the neuron, including terminals whose partners belong to small segments that are not connected to any proofread neuron ("twigs";[2]). **B** Mean SI values calculated using the Princeton synapse detector[3] and the Buhmann synapse detector [1]. Neurons were binned according to their SI value from the Princeton detector, and circle size indicates the number of neurons in each bin. **C** Same as in B, shown separately for each superclass.

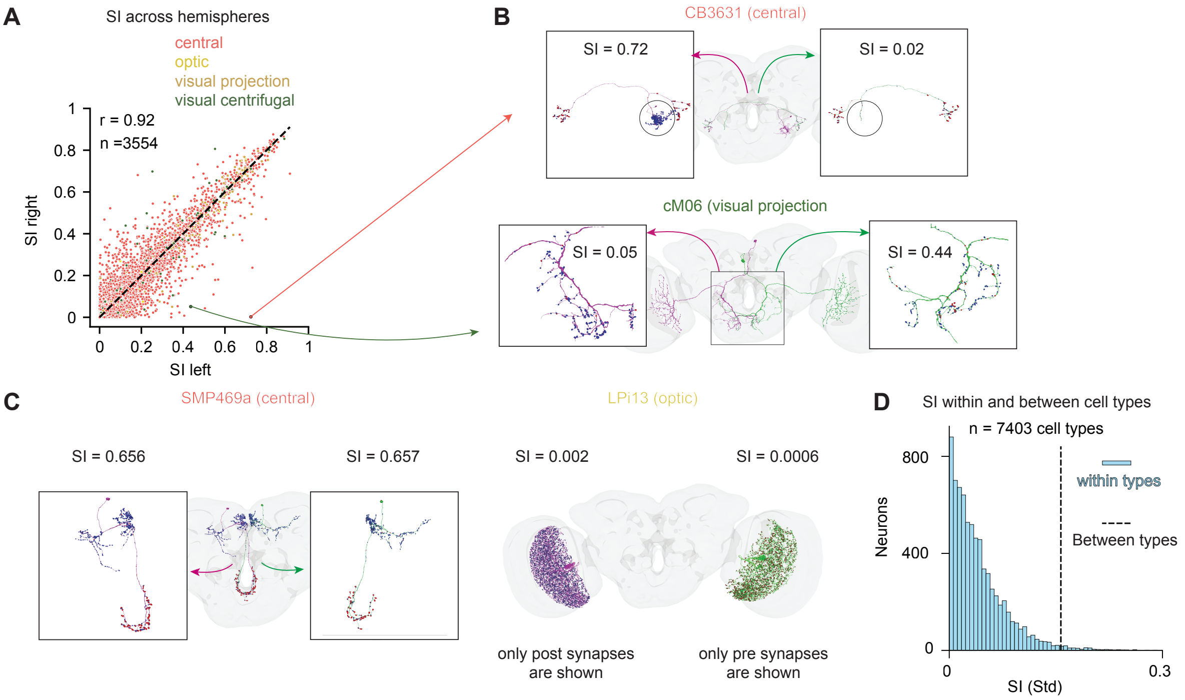

**Supp Fig. 2 – S2. A** SI values of homologous left–right neuron pairs belonging to the same primary type variant (n = 3,554 cell types containing 1 cell per hemisphere). Each point corresponds to one pair, with the SI of the left-hemisphere neuron plotted on the x-axis and the SI of the right-hemisphere neuron plotted on the y-axis (Pearson r = 0.92). **B** Two example neuron pairs with low left–right SI symmetry. Top: central brain CB3631 neurons with asymmetric SI values arising from the presence of a prominent postsynaptic branch in one hemisphere that is reduced or absent in its homolog. Bottom: visual centrifugal cM06 neurons with SI differences driven by unequal distributions of presynaptic sites between hemispheres. Arrow tails indicate the corresponding points in panel A. **C** Two example neuron pairs with high left–right SI symmetry: SMP469a (central) and LPi13 (optic), with high and low SI values in both hemispheres, respectively. For LPi13, only postsynaptic sites are shown for the left neuron and only presynaptic sites for the right neuron because of the high synapse density. **D** Histogram of within-type SI standard deviations (SD) across 7,403 primary neuron types (≥2 neurons per type); the dashed line indicates the between-type SD, calculated from the mean SI value of each type. Welch's ANOVA on SI across primary types revealed a strong effect ($F(7402, 3972.97) = 67536.91$, $p < 0.001$, $\eta^2 p = 0.826$) , indicating that between-type SI variability substantially exceeds within-type SI variability.

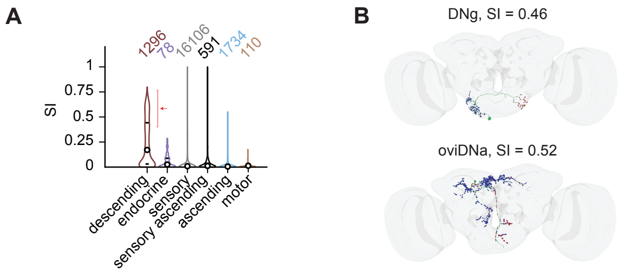


**Supp Fig. 2 - S3 A** Violin plots showing SI distributions across the non-intrinsic neuron superclasses. Dots mark medians and vertical lines show the interquartile range; population sizes are indicated above each violin. Red arrow indicates high SI values for descending neurons, despite most of their outputs being absent from the dataset. **B** Two examples of descending neurons with high SI values (from the SI range highlighted by the red arrow in panel A): DNg (SI = 0.46) and oviDNa (SI = 0.52).

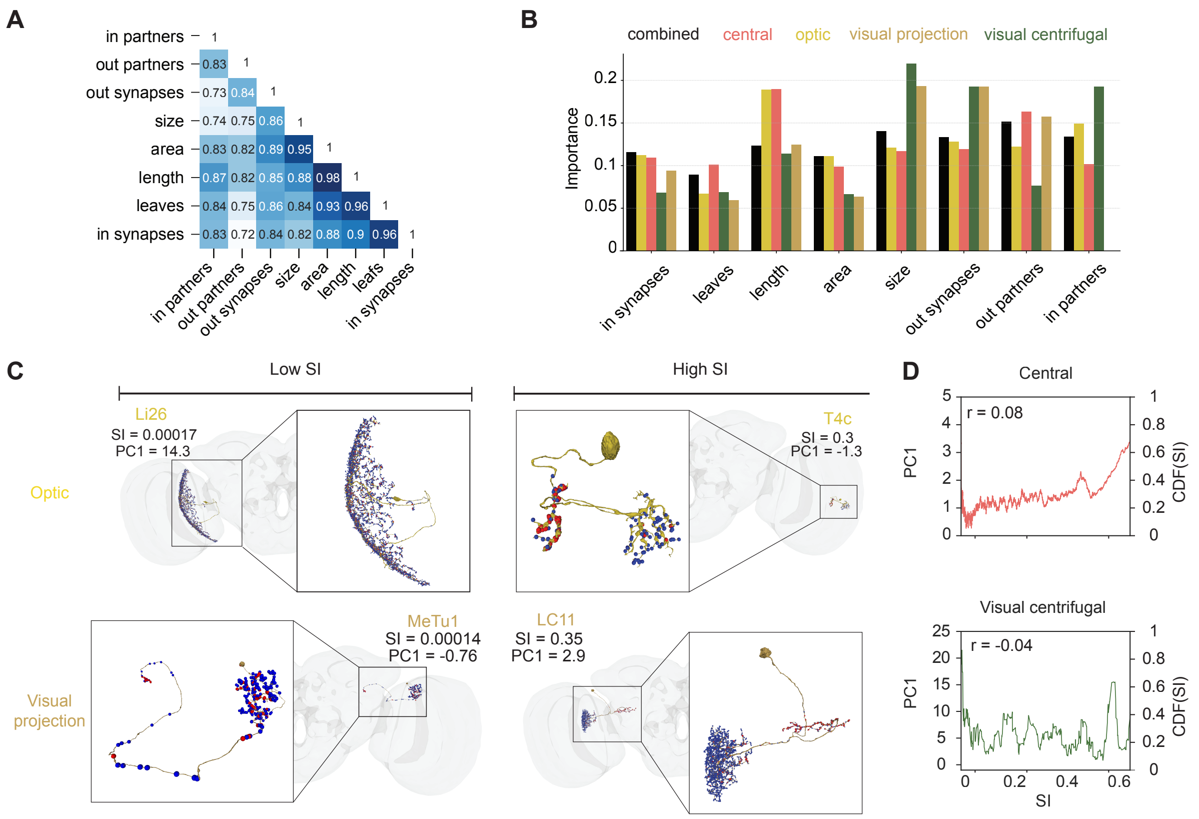


**Supp Fig. 2 - S4 A** Lower-triangular correlation matrix showing pairwise correlations among the eight features used for SI prediction. **B** Feature importance values from the random forest model used in Fig. 2E to predict Segregation Index (SI), shown separately for neurons in each superclass (see Methods). **C** Example optic (top) and visual projection (bottom) neurons. For optic neurons, 'simple' cells are more polarized, while the opposite is true for visual projection neurons (Fig. 2G). **D** Relationship between SI and the first principal component (PC1) in central (top) and visual centrifugal (bottom) neurons. Solid colored lines show the rolling mean of PC1 as a function of SI. A weak positive correlation between PC1 and SI is observed in central neurons ($r = 0.08$; $P < 0.0001$), whereas no significant association is detected in visual centrifugal neurons ($r = -0.04$; $P = 0.33$).

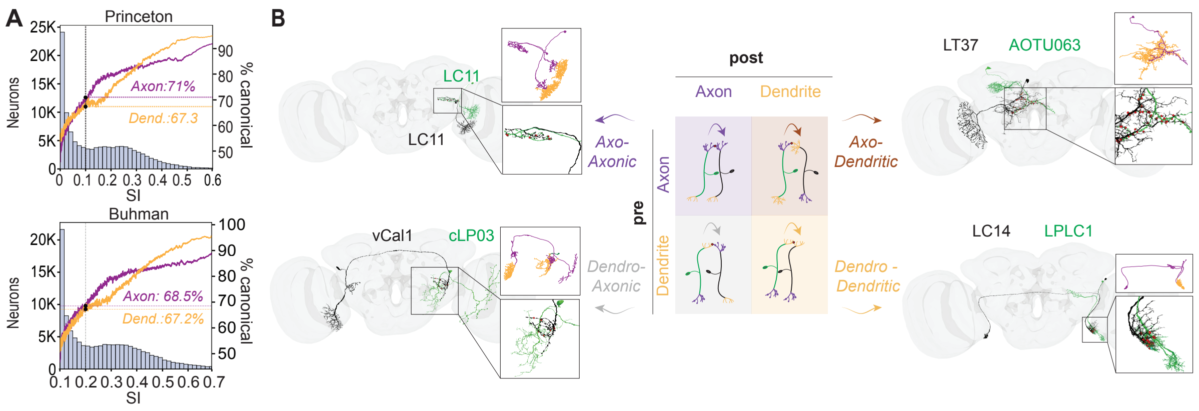


**Supp Fig. 3 - S1 A** Fraction of canonical synaptic terminals: postsynaptic terminals on dendrites, presynaptic terminals on axons, as a function of the SI (rolling mean with window = 400 for sorted SI). Using Princeton synapses, for SI = 0.1 (dashed lines), 71% of the synaptic terminals on axons are presynaptic and 67.3% of the terminals on dendrites are postsynaptic compared to 68.2% and 67.2%

using the Buhmann detector. Those values (using Princeton synapses) are used as thresholds to determine whether a given compartment (dendrite or axon of a single neuron) is considered ‘mixed’ (under the threshold) or not (equal to or above the threshold). For example, with an SI threshold of 0.1, an axonal compartment in which 80% of terminals are presynaptic is classified as non-mixed, whereas a compartment with only 60% presynaptic terminals is classified as mixed. **B** Schematics and examples for the four possible connection types: axon-to-dendrite (AD), axon-to-axon (AA), dendrite-to-dendrite (DD), and dendrite-to-axon (DA). Each configuration is illustrated with an example pair of neurons (presynaptic neuron in green and postsynaptic neuron in black). Axonal compartments are shown in purple and dendritic compartments in orange, indicating the specific subcellular regions where synaptic contacts occur. See links for Neuroglancer in Table 2.

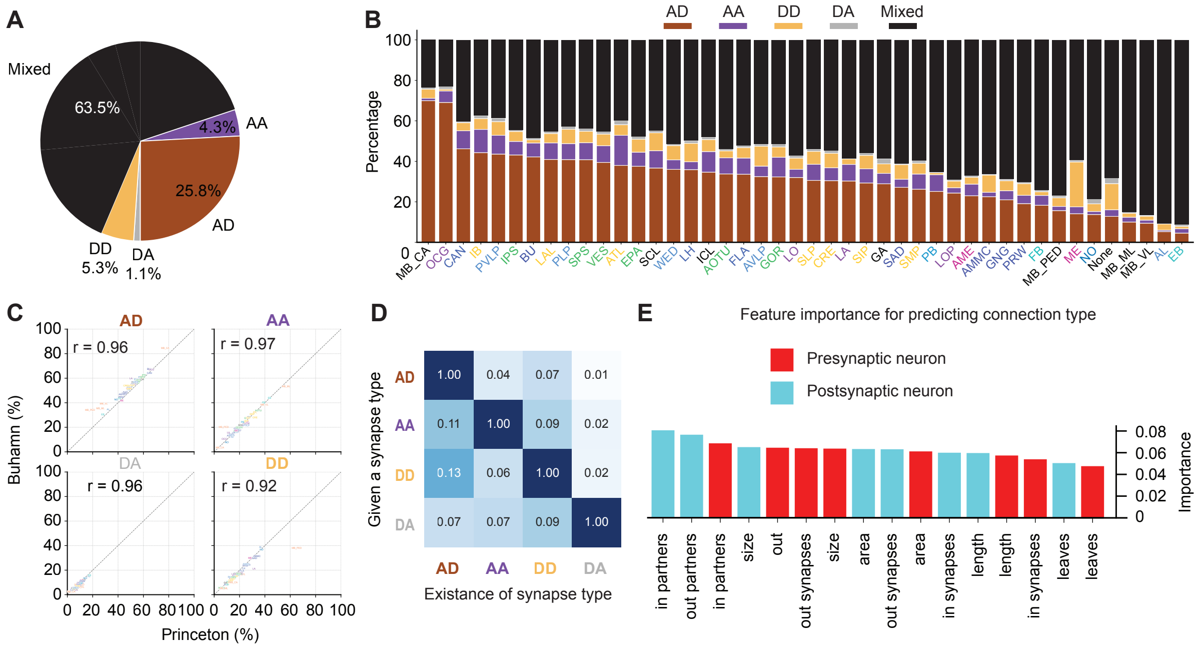


**Supp Fig. 3 - S2.** **A** Percentages of canonical, non-canonical, and mixed synapses across intrinsic neuron superclasses, using the Buhmann synapse dataset (44,342,339 synapses). **B** Percentages of synaptic connection types across neuropils, using the Buhmann synapse detector. **C** Comparison of synaptic-type proportions estimated using the Princeton and Buhmann synapse detectors, shown separately for each synapse type. Pearson correlation coefficient is shown for each type. **D** Conditional probabilities of synaptic-type co-occurrence, conditioned on the presence of a given connection type. For example, among all neuron pairs connected by at least one AD synapse (first row), 4% also contain at least one AA synapse. **E** Feature importance values for the random forest model using eight presynaptic and eight postsynaptic features to predict synapse type (see model performance in Fig. 3J and Methods). Higher values indicate greater influence of the feature on model predictions (see Methods).

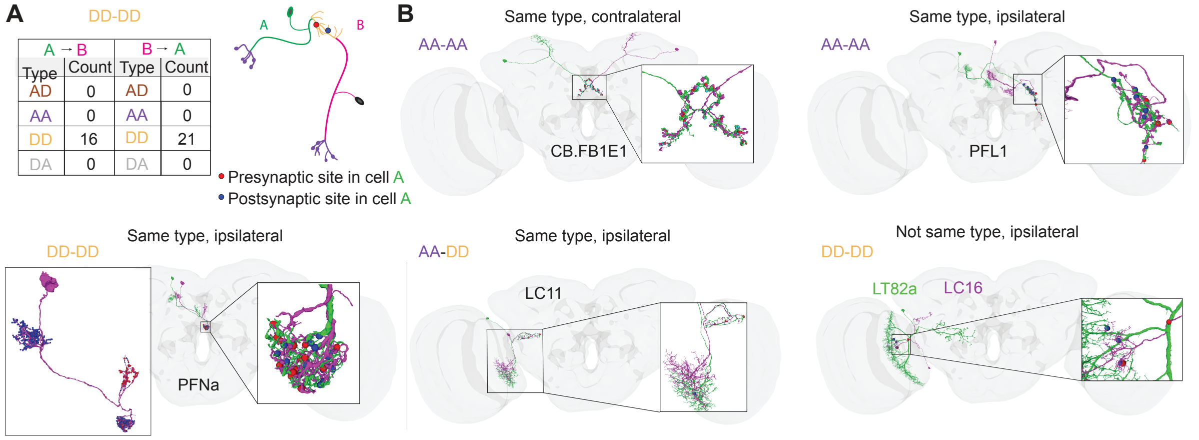


**Supp Fig. 5–S1. A** Example of a reciprocal DD↔DD connection between two central complex neurons of the same cell type (PFNa). There are 16 and 21 DD synapses in the two directions of the connection. Inset: distribution of all presynaptic (red) and postsynaptic (blue) terminals for the purple PFNa neuron. This neuron contains two regions dominated by postsynaptic terminals, one of which forms reciprocal connections with the green PFNa neuron shown in the example (see also Supp Fig. 6 - S2). **B** Example neuron pairs illustrating AA↔AA (the central complex neurons FB1D and PFL1), DD↔DD (visual projection neurons LT82a and LC16), and AA↔DD (visual projection neurons of type LC11) reciprocal motifs.

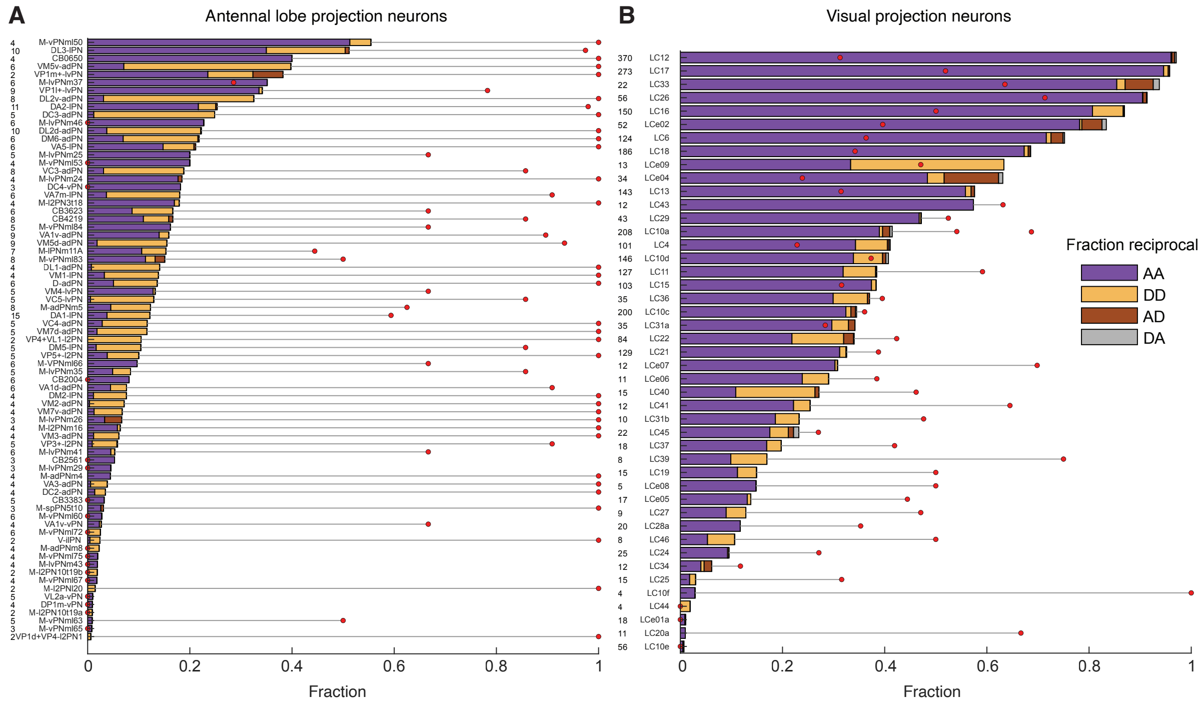


**Supp Fig. 6–S1. A** Bar plot showing the distribution of synapse types connecting neurons within the same cell type for antennal lobe projection neurons (ALPNs). Red circles indicate, for each cell type, the fraction of connected pairs that are reciprocally connected. Left: number of neurons belonging to each cell type. **B** Same as in A for visual projection neurons.

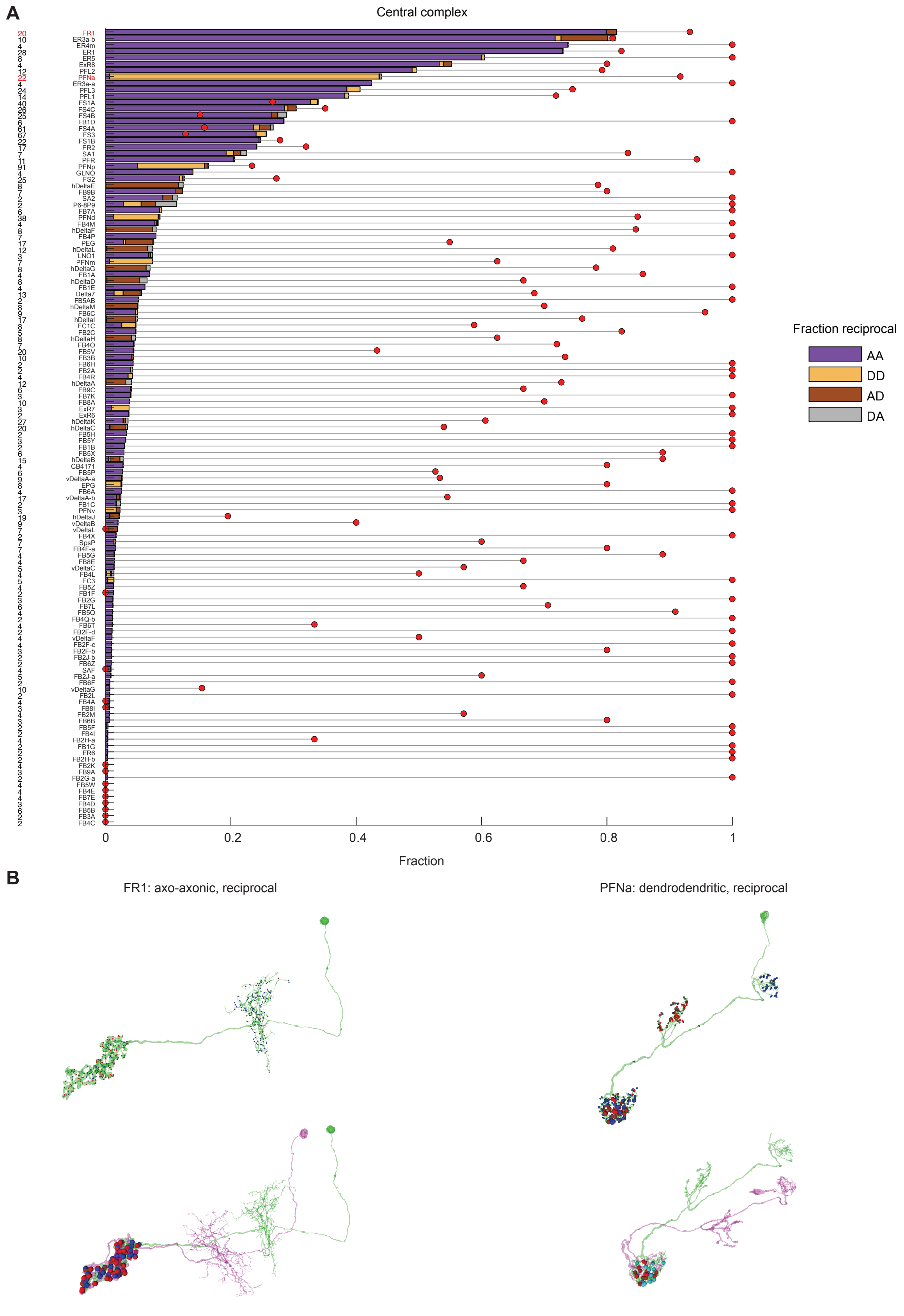


**Supp Fig. 6–S2. A** Bar plot showing the distribution of synapse types connecting neurons within the same cell type for antennal lobe projection neurons (ALPNs). Red circles indicate, for each cell type, the fraction of connected pairs that are reciprocally connected. Left: number of neurons per cell type.

**B** Examples of reciprocal connections between pairs of neurons of the same cell type. Left: cell type FR1; right: cell type PFNa. For each example, the top panel shows all presynaptic and postsynaptic terminals of one neuron (red and blue, respectively). The bottom panel shows both neurons in the pair, with only the synapses connecting the two neurons displayed. Red and blue markers indicate presynaptic and postsynaptic terminals, respectively, from the perspective of the green neuron.

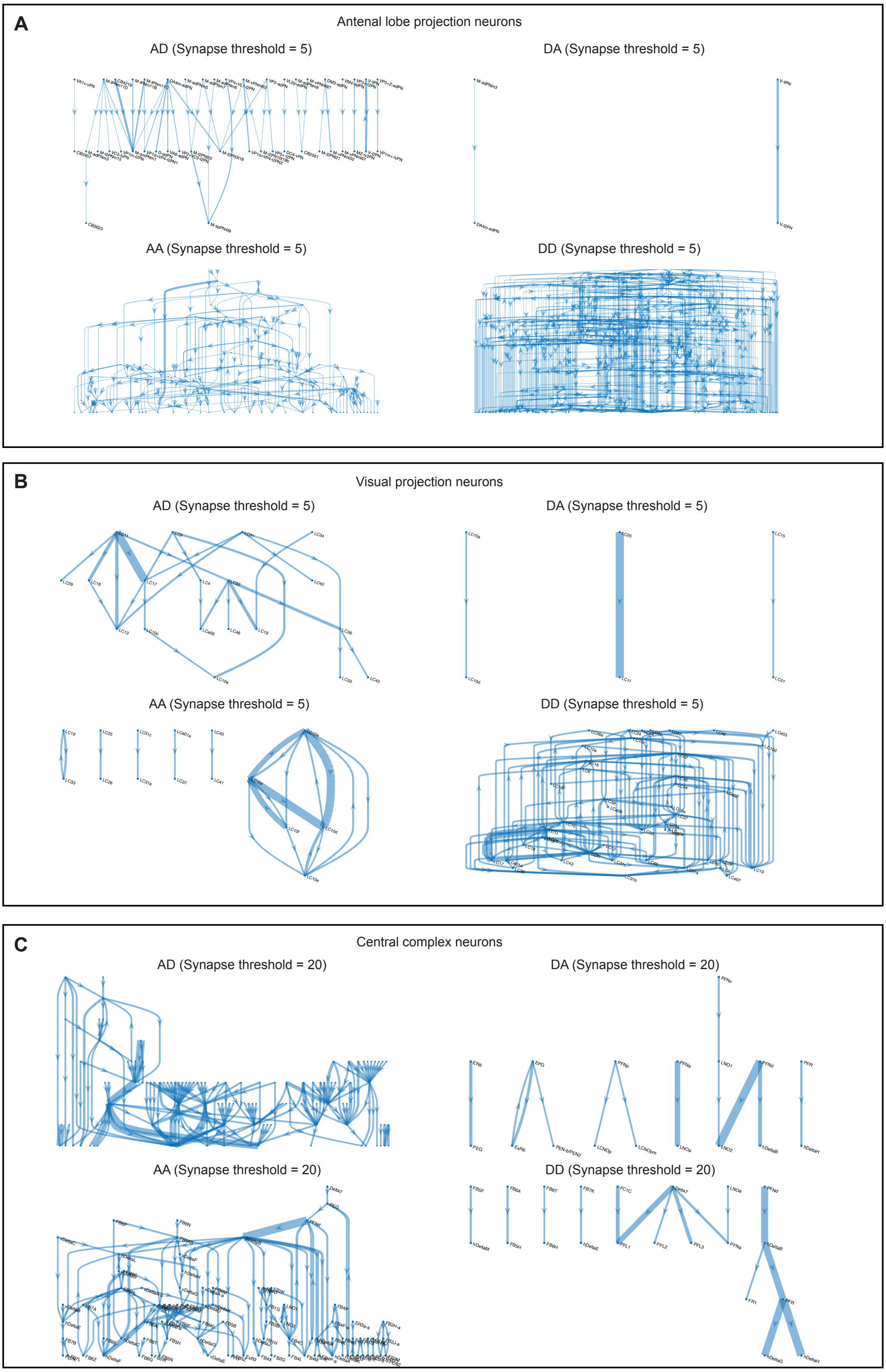


**Supp Fig. 6 – S3**. Directed connectivity between cell types for three neuronal populations—antennal lobe projection neurons (ALPNs), visual projection neurons, and central complex neurons - shown separately for each connection type (AD, AA, DD, and DA). Only connections between neurons belonging to different cell types are included. Each node represents a single cell type. Arrow width scales with the number of synapses. For clarity, only connections exceeding a minimum synapse threshold are shown. A threshold of 5 synapses was used for ALPNs and visual projection neurons, and a higher threshold of 20 synapses was used for central complex neurons. DD connections dominate among ALPNs and visual projection neurons, whereas AD and AA connections are more prominent among central complex neurons.

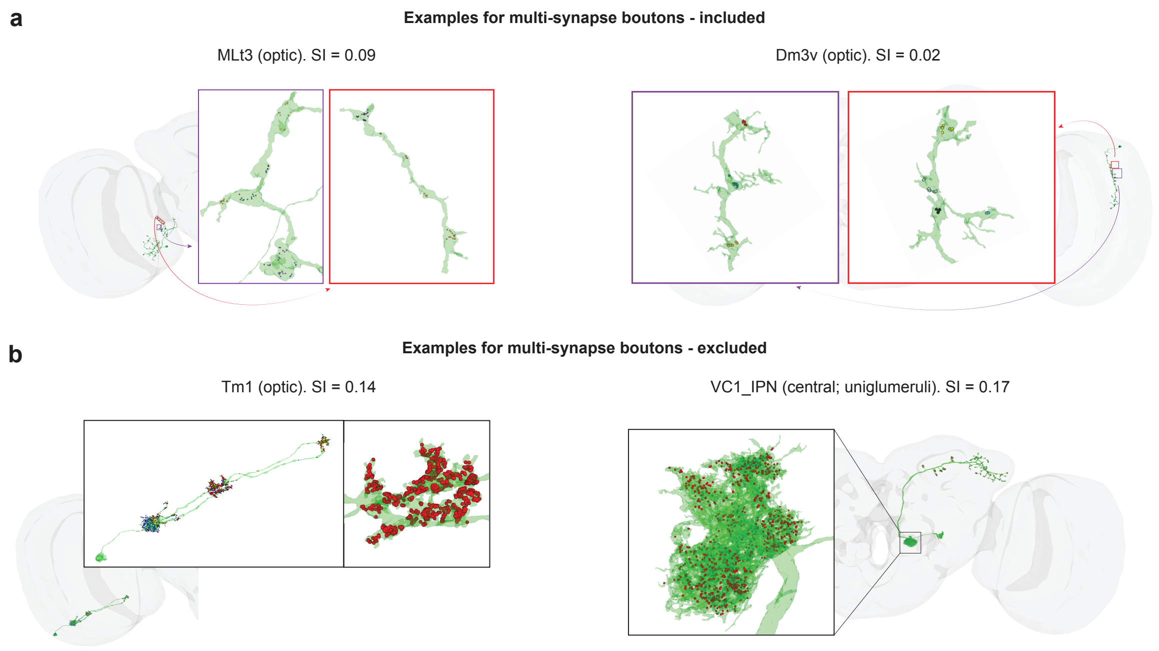


**Supp Fig. 7–S1. A** Examples of boutons in two neurons. Each color represents a distinct presynaptic cluster corresponding to an individual multisynaptic bouton (MSB). Left: example neuron of cell type MLt3. Right: example neuron of cell type Dm3v. **B** Examples of neurons lacking typical bouton organization. Left: an optic neuron of cell type Tm1 showing dense presynaptic regions that cannot be reliably separated into distinct spatially segregated MSBs. Right: an example uniglomerular neuron exhibiting a single large, dense presynaptic region that does not correspond to a single MSB. Links to FlyWire examples are provided in Table 2. The list of cell types excluded from MSB analyses is detailed in Table 4.

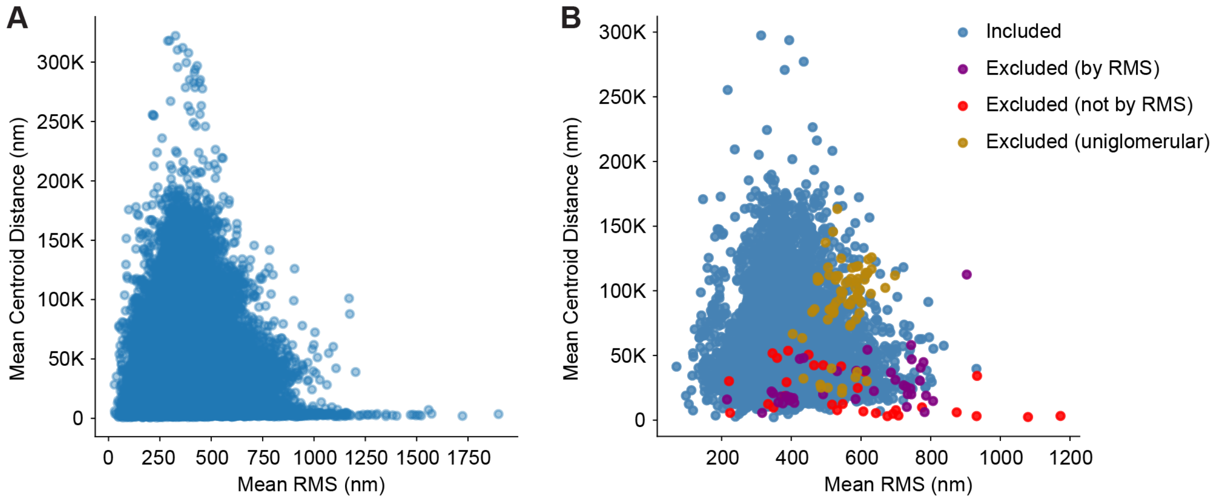


**Supp Fig. 7–S2. A** Mean RMS (root mean square distance from presynaptic terminals to their cluster centroid in nanometers) and mean distance between cluster centroids (in nanometers), calculated for each neuron in FlyWire/FAFB. **B** Mean RMS and mean centroid distance averaged per cell type (each dot represents a single cell type). Several cell types were excluded from analyses that relied on automated MSB detection. These included all uniglomerular neurons (yellow; see Supp Fig. 7–S1), cell types that include cells with high mean RMS values (purple; Mean RMS > 850 in panel A), indicating large and spatially dispersed clusters, and additional cell types that were manually excluded (red; see Methods and Table 4).

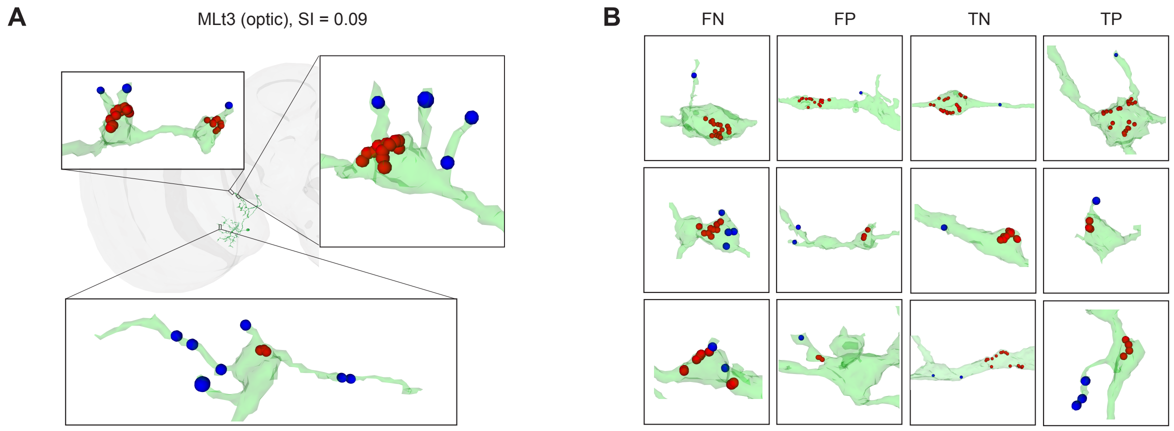


**Supp Fig. 7–S3. A** Examples of postsynaptic terminals on MSBs, including cases with one or multiple postsynaptic terminals located on filopodia extending from MSBs (cell type MLt3). **B** Examples from each category of the confusion matrix shown in Fig. 7D. Three representative examples are shown for each category (FN, FP, TN, and TP), illustrating the performance of the detector in identifying postsynaptic terminals on MSBs (either directly on the surface of the MSB or on filopodia extending from the MSB). To provide a more stringent evaluation (see Methods), the pseudorandom negative examples were restricted to contain at least one postsynaptic terminal near the MSB. Consequently, postsynaptic terminals are visible in all examples. For example, in the three FN (false negative) examples, the detector classified the MSB as negative (no adjacent postsynaptic terminals), whereas one or more postsynaptic terminals are present on the MSB, making the prediction false.

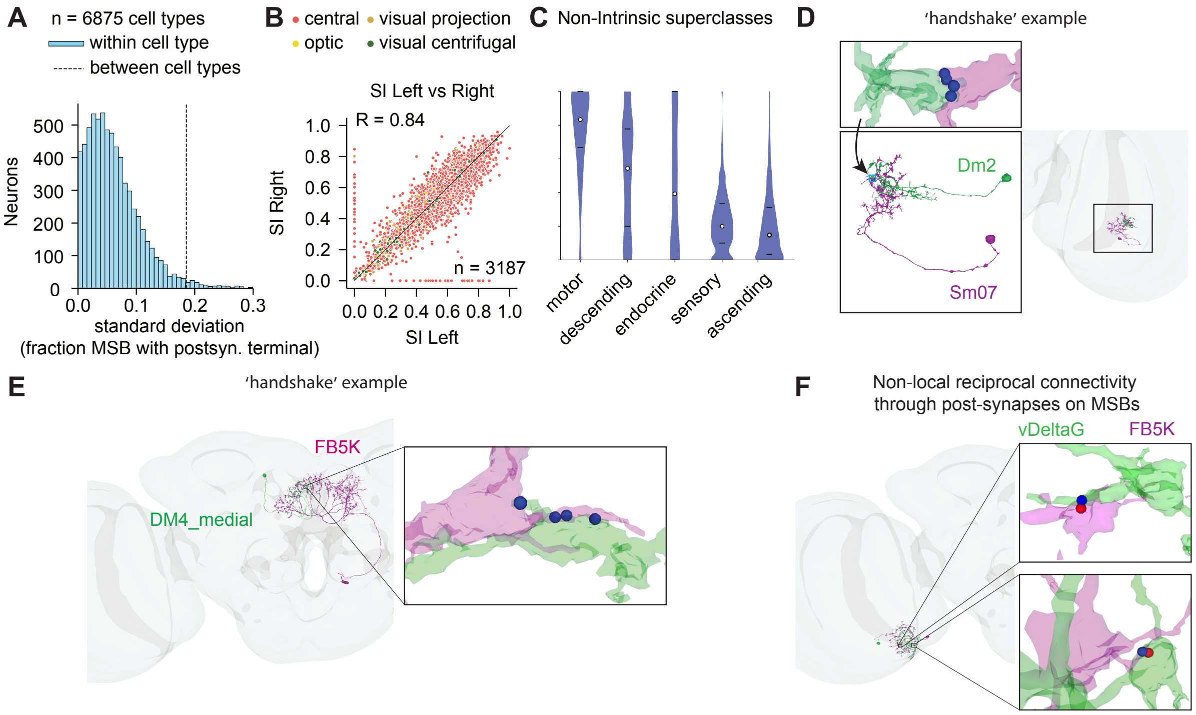


**Supp Fig. 7–S4. A** For each neuron belonging to an annotated cell type with two or more neurons in the dataset, the fraction of MSBs containing one or more postsynaptic terminals was calculated. For each cell type $i$, two quantities were then computed: $S_i$, the standard deviation of this fraction across neurons of that type, and $M_i$, the corresponding mean. The histogram ("within cell type") shows the distribution of $S_i$ across all cell types ($N = 6{,}875$ cell types with ≥2 neurons per type), whereas the dashed vertical line ("between cell types") indicates the standard deviation across the mean values $M_i$. In a random null model, between-type variability is expected to be smaller than the mean within-type variability. **B** Left–right correlation of the fraction of MSBs containing one or more postsynaptic terminals (n = 3,491 cell types with exactly one neuron per hemisphere). Points are colored by superclass; the dashed line indicates the identity line (x = y). Pearson r = 0.84. **C** Fraction of MSBs containing one or more postsynaptic terminals for non-intrinsic neurons, separated by superclass and neuronal compartment, for neurons with SI ≥ 0.1. The fraction was first calculated per neuron. White dots indicate medians and black lines indicate quartiles (similar to Fig. 7F, where the fraction was calculated for intrinsic neurons). **D** Two closely positioned MSBs belonging to different neurons, each containing postsynaptic terminals. Postsynaptic terminals are shown as blue dots: the two upper terminals belong to the purple neuron and the two lower terminals belong to the green neuron, illustrating a local reciprocal connection (presynaptic terminals are omitted for clarity). **E** Same configuration as in D except that the upper-left postsynaptic terminal (blue) belongs to the green neuron, whereas the remaining three postsynaptic terminals belong to the purple neuron. **F** Example of a reciprocal connection between two neurons that is not locally confined to adjacent MSBs (and therefore is not classified as a handshake). Each MSB belongs to a different neuron and is located in a separate spatial region, yet the two neurons are connected bidirectionally through postsynaptic terminals on MSBs. Top: postsynaptic terminal from the green neuron (blue) and an MSB of the purple neuron (presynaptic terminals in red). Bottom: the reciprocal connection, in which the presynaptic terminal is located on the green neuron. This example corresponds to the yellow category in Fig. 7K, which includes reciprocal connections involving postsynaptic terminals on MSBs in one or both directions, provided that the two neurons are reciprocally connected.

**References**

Buhmann, J. *et al.* Automatic detection of synaptic partners in a whole-brain Drosophila electron microscopy data set. *Nat. Methods* **18**, 771–774 (2021). 1.

Dorkenwald, S. *et al.* Neuronal wiring diagram of an adult brain. *Nature* **634**, 124–138 (2024). 2.

Yu, S.-C. *et al.* New synapse detection in the whole-brain connectome of *Drosophila*. *bioRxiv* (2025) doi:10.1101/2025.07.11.664377. 3.
